# Diurnal rhythmicity in metabolism and salivary effector expression shapes aphid performance on host plants

**DOI:** 10.1101/2024.01.20.576473

**Authors:** Jinlong Han, Daniel Kunk, Meihua Cui, Yoshiahu Goldstein, Vered Tzin, Vamsi J. Nalam

## Abstract

Diurnal rhythms influence insect behavior, physiology, and metabolism, optimizing their performance by adapting to daily changes in the environment. While their impact on agricultural pests has been briefly explored, our understanding of how these rhythms drive adaptative responses in pest biology and influence host colonization remains elusive. Here, we show that a notorious global aphid pest, *Rhopalosiphum padi*, exhibits distinct diurnal patterns in feeding behavior, with elevated honeydew excretion at night and extended phloem salivation during early nighttime. Temporal aphid transcriptome profiling reveals four diurnally rhythmic clusters, two of which peak at night, exhibiting enrichment in carbohydrate and amino acid metabolism. Beyond the established role in manipulating host responses and allowing sustained feeding, our study reveals novel evidence of cyclical fluctuations in salivary effector expression in an insect species. Silencing key effector genes, peaking in expression during the increased nighttime salivation, results in a more pronounced reduction in aphid excretion activity on host plants during the night compared to the day, a phenomenon not observed on artificial diets. A better understanding of aphid diurnal rhythms and their roles in shaping aphid performance provides a promising avenue to refine and optimize pest management, granting a strategic advantage for minimizing crop damage.

## Introduction

The recurring cycles of day and night caused by Earth’s daily rotation profoundly influence all life, including plants and their pests. All living organisms have evolved internal biological clocks (i.e., circadian clocks) to anticipate and respond to cyclical changes in their environments by orchestrating various behavioral and physiological processes ^1^. Diurnal and circadian rhythms entail daily biological cycles, with circadian rhythms representing a subset characterized by internally sustained oscillations that persist in the absence of external cues ^2^. Our understanding of these rhythms has led to profound insights into mechanisms governing the intricate biological processes that shape various daily and seasonal behavioral patterns in diverse taxa, including insects and plants. Despite a significant body of knowledge on diurnal rhythms in model organisms and host-parasite interactions ^3,4^, research on agricultural pests and their interactions with plant hosts remains underexplored. Given the significant threat pests pose to crop production, deciphering the temporal aspects of their behavior, physiology, and gene expression is a crucial step toward developing more effective pest control strategies ^5,6^.

Aphids are among the most devastating pests worldwide, causing significant damage to various economically important crops through their consumption of sugar-rich phloem sap, delivery of phytotoxic salivary secretions, transmission of plant-pathogenic viruses, and promoting the growth of sooty-mold on their sugar-rich excrement ^7^. Studies on circadian clock-regulated plant defenses against arthropod pests reveal that pests strategically anticipate and exploit plant rhythms. For example, the cabbage looper (*Trichoplusia ni*), a generalist herbivore, shows rhythms in feeding, with peak feeding occurring during the latter part of the day ^8^. Conversely, cell-content feeders like spider mites (*Tetranychus urticae*) exploit a nighttime “lull” in jasmonate-induced defenses, causing more damage during the night ^9^. Given that aphids rely entirely on acquiring phloem sap from plants for their reproduction and survival, they likely need to anticipate, respond to, and counteract the challenges posed by daily rhythms in their hosts. Notably, green peach aphids (*Myzus persicae*) feeding on Arabidopsis plants exhibit peak honeydew droplet excretion during the day ^10^. These diurnal patterns in aphids extend beyond honeydew excretion, encompassing aspects such as locomotion, larviposition, sex hormone release, egg hatching, and feeding ^11–16^. While these studies have begun to broaden our understanding of aphid diurnal rhythms, our knowledge about how aphids regulate their rhythmic behaviors and interact with their hosts over daily cycles is lacking.

The high energy demand for aphid reproduction and the ‘nitrogen and sugar barrier’ in phloem sap necessitate aphids to feed incessantly from their host plants ^17,18^. To facilitate this continuous feeding, aphids synthesize watery saliva in their salivary glands and secrete it into plant cells. This watery saliva comprises a complex blend of proteins with enzymatic activity, including cell-wall degrading enzymes, proteases, polyphenol oxidases, oxidoreductases, and peroxidases, as well as diverse calcium-binding proteins, and many others with unknown functions ^19^. Collectively termed effectors, these salivary proteins modify plant defense and physiology to better facilitate aphid feeding, survival, and reproduction ^20^. With the advancement in omics technologies, numerous salivary effector candidates have been identified in several aphid species ^21–23^. An early and renowned example is C002, a protein with an unknown function that is essential for continuous feeding from sieve elements ^24^. Silencing C002 expression results in lethal outcomes in the pea aphid, *Acyrthosiphon pisum*, and diminished fecundity in the green peach aphid, *Myzus persicae* ^25,26^. Despite the pivotal roles of salivary effectors in the aphid colonization of host plants, knowledge about whether their expression or function undergoes diurnal rhythms is unknown.

Another consequence of a sugar-rich diet is that aphids must process and excrete excess sugar to maintain an appropriate osmotic balance ^18^. Aphid osmoregulation involves physiological processes such as the breakdown of sucrose into glucose and fructose, their polymerization and expulsion as honeydew, and the rapid cycling of water between the midgut and hindgut ^27,28^. Additionally, aphids feed during the night when phloem sugar content is lower ^16^ and occasionally drink dilute sap from the xylem, a behavior hypothesized to contribute to rehydration ^29^. At the molecular level, several genes involved in osmoregulation have been identified in aphids, and their silencing has been shown to elevate osmotic pressure in aphid hemolymph and impair performance ^30–32^. A prior study also showed periodic variations in the expression of osmoregulatory genes throughout a 24 h period in the bird cherry-oat aphid, *Rhopalosiphum padi* ^16^. Although diurnal fluctuation in phloem sap composition and its impact on aphid performance has been reported ^33–36^, an in-depth investigation into diurnal variation of aphid osmoregulation at the behavioral and molecular level is currently lacking.

Here, our study shows the diurnal rhythmicity of feeding behavior and gene expression in *R. padi*, a serious global pest of cereals, and its implications for aphid-host interactions ^37^. We demonstrate diurnal variations in honeydew excretion and specific feeding activities, notably the salivation process into the phloem. Our temporal analysis of the aphid transcriptome reveals diurnally rhythmic gene clusters with unique expression patterns and pathway enrichment, rationalizing the observed diurnal changes in our aphid feeding assays. Significantly, diurnally rhythmic putative salivary effectors were identified, with many showing peak expression during the increased nighttime salivation. By integrating behavioral and molecular experiments, we unveil a significant time-oriented role of osmoregulatory and putative salivary effector genes in aphids feeding on host plants. These findings provide novel insights into the diurnal regulation of aphid behavior, metabolism, and salivary effectors, highlighting their crucial roles in the plant-aphid relationship.

## Results

### Diurnal rhythmicity in *R. padi* feeding behaviors

Diurnal variation of honeydew excretion in *R. padi* was investigated under a 12L:12D cycle for 48 h on wheat plants. Our study found that the number of honeydew droplets produced per aphid was higher during nighttime (12.09 ± 0.75, mean ± standard error of mean) as opposed to daytime (8.42 ± 0.57) (Fig. 1a). The experiment was repeated on an artificial diet to determine if the observed patterns were independent of plant-derived cues. Although fewer honeydew droplets were produced on the diet overall, a higher number of honeydew droplets was consistently produced per aphid on the diet during the night (0.23 ± 0.05) compared to the day (0.09 ± 0.03) (Fig. 1b).

**Figure 1.**
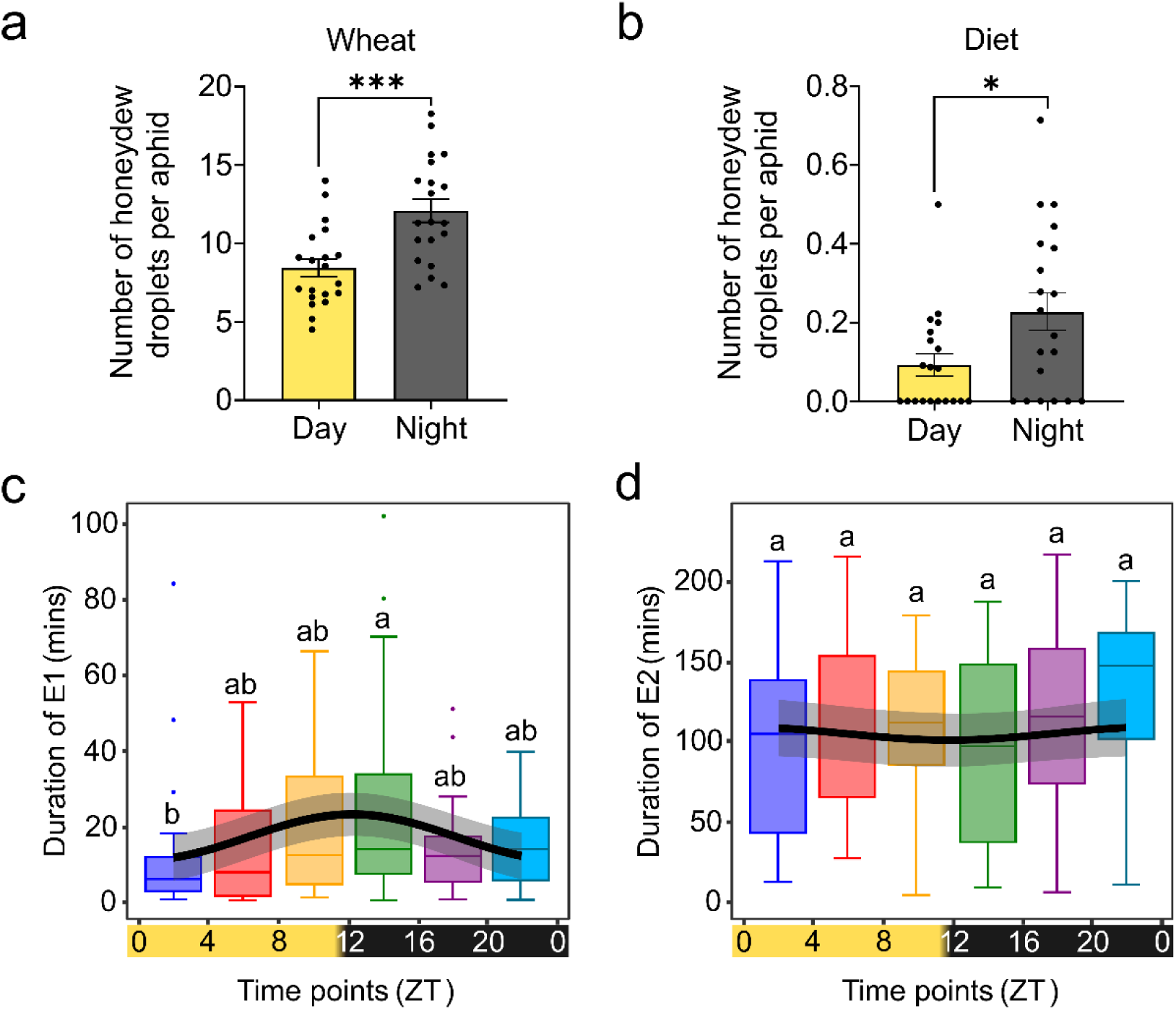
Diurnal rhythmicity of aphid feeding behaviors. **a, b** The number of honeydew droplets produced by aphids was measured at the end of each 12h day and night cycle for 48h. Average number (±SE) of honeydew droplets produced per aphid on wheat plants (**a**, n=10) and artificial diet (**b**, n=10). Significant differences between day and night were tested by the two-tailed Mann-Whitney test (wheat: \*\*\**p=*0.0006 and diet: \**p=*0.0411). **c, d** Aphid feeding activity was monitored every 4h over a 24h period, 0zt (n=25), 4zt (n=21), 8zt (n=26), 12zt (n=23), 16zt (n=29), and 20zt (n=16). The total time (±SE) spent by aphids in phloem salivation (E1) and phloem ingestion (E2) are shown in (**c**) and (**d**), respectively. Solid lines, boundaries, and whiskers indicate the median, 25^th^, and 75^th^ percentiles and 1.5 interquartile ranges, respectively. Points indicate outliers. Different letters above the boxes denote significant differences (Kruskal-Wallis test followed by Dunn’s multiple comparisons adjusted by Bonferroni for *p*-values, adj. *p*<0.05). Diurnal rhythmicity of E1 (**c**) and E2 (**d**) waveforms over 24h was assessed with Cosinor model. The curve line and shaded area represent the fitted periodic sinusoidal curve and 95% confidence band, respectively. Significance is only established for a diurnal rhythm if the Cosinor *p*-value is <0.05 (E1: *p*=0.0119 and E2: *p*=0.7313). Yellow and black bars at the x-axis mirror the day and night. ZT = Zeitgeber time. Source data and full statistical summary are provided in the Source Data file.

The diurnal feeding behavior of *R. padi* was further elucidated using the electrical penetration graph (EPG) technique by measuring feeding at 4 h intervals over a 24 h period (12L:12D). Significant differences were observed in various aphid feeding activities over each timepoint (Supplementary Data 1), including the phloem salivation (E1) phase, which represents the injection of saliva into plant sieve elements. Adult *R. padi* spent the longest time in E1 between 12-16 zeitgeber time (zt, refers to an environmental agent or event that provides the stimulus to set or reset a biological clock) and the shortest time between 0-4zt, each of which represents the time period directly following the removal or introduction of the light cues, respectively (Fig. 1c). We also identified a diurnal rhythm in the total duration of E1, with an acrophase (the estimated time in a rhythmic cycle when a peak or maximum value occurs) occurring at 9.8zt (Fig. 1c, Supplementary Table 1), suggesting a potential link between light transitions and the initiation of salivation. Interestingly, despite the higher honeydew excretion at night (Fig. 1a), the total time spent in phloem ingestion (E2 phase) did not differ over the 24 h period. No diurnal rhythm was detected for the E2 phase (Fig. 1d). Therefore, we hypothesize that the observed day-night variations in salivation and honeydew excretion are associated with diurnally regulated genes involved in the expression of salivary effectors and metabolic pathways linked to aphid feeding, such as osmoregulation.

### Temporal gene expression patterns in *R. padi* reveal diurnal rhythmicity

Temporal transcriptome profiles were collected from *R. padi* whole bodies over a 24 h period to identify diurnal rhythms in gene expression. A total of 49,486 transcripts were assembled with an average sequence length of 2,204 bp. The overall read mapping rate against the *R. padi* reference genome and Benchmarking Universal Single-Copy Orthologs (BUSCO) score for complete sequences were 97% and 99%, respectively (Supplementary Table 2). Of the total assembled transcripts, 4,460 (9.2%) showed significant diurnal rhythmicity over a 24 h period (Supplementary Data 2). 2,913 and 1,547 rhythmic transcripts were identified to have acrophases during the day (0-12zt) and night (12-24zt), respectively. Most transcripts displayed acrophases between 4-8zt and 17-19zt, representing midday and midnight periods, respectively (Fig. 2a).

**Figure 2.**
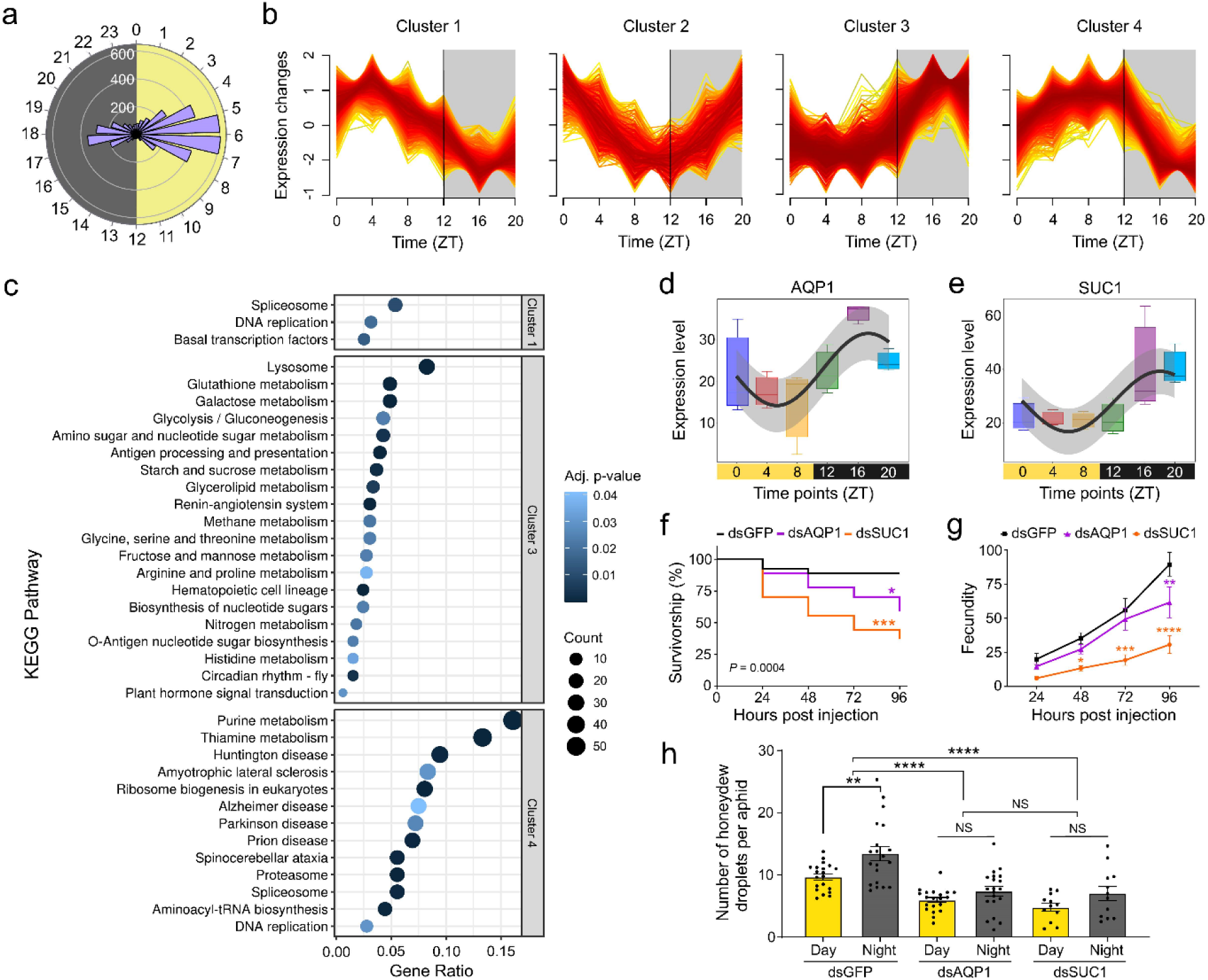
Diurnal rhythmicity of gene expression in *R. padi* and their roles in aphid feeding. **a** Acrophase frequency of diurnally rhythmic transcripts in *R. padi*. Each bar represents the number of rhythmic sequences with acrophase values in each 24h period. Numbers around the outer sphere indicate zeitgeber time (ZT). **b** Soft-clustering of diurnally rhythmic transcripts. High, medium, and low membership scores are color-coded with red, yellow, and blue lines. **c** Top-10 enriched KEGG pathways for diurnally rhythmic transcripts in each cluster. Circle size and color represent enriched gene number in a pathway, and Benjamini-Hochberg adjusted *p*-value, respectively. Gene ratio indicates the percentage of diurnally rhythmic transcripts of a cluster. **d, e** Expression profile of two diurnally rhythmic osmoregulatory genes, aquaporin 1 (AQP1, **d**) and gut sucrase 1 (SUC1, **e**). Solid lines, boundaries, and whiskers of each box indicate median, 25^th^, and 75^th^ percentiles and 1.5 interquartile ranges. Diurnal rhythmicity of AQP1 (*p*=0.031) and SUC1 (*p*=0.013) was determined with the Cosinor model. The curved line and shaded area represent the fitted periodic sinusoidal curve and 95% confidence band, respectively. **f** Kaplan-Meier survival curves and **g** fecundities of aphids after treatment with dsRNA AQP1 (dsAQP1, n=3), dsRNA SUC1 (dsSUC1, n=3), and dsRNA GFP (dsGFP, n=3, negative control). Each replicate (*n*) contains nine individual aphids. Significant differences in survivorship and fecundity were tested using a log-rank test with Benjamini-Hochberg adjustment (dsAQP1 vs dsGFP: \**p*=0.0272 and dsSUC1 vs dsGFP: \*\*\**p*=0.0003) and a repeated measures ANOVA with Dunn’s multiple comparisons test (\**p*<0.05, \*\**p*<0.01, \*\*\**p*<0.001, \*\*\*\**p*<0.0001), respectively. **h** Average number (±SE) of honeydew droplets produced per aphid after treatment with dsAQP1 (n=5), dsSUC1 (n=3), and dsGFP (n=5) on wheat. Each replicate (n) contains five individual aphids. Yellow and black bars represent the average number of droplets produced at the end of each 12:12h day and night for 4 days (Mann-Whitney test, \*\**p*<0.01, \*\*\*\**p*<0.0001, NS=no significant difference). Source data and full statistical summary are provided in the Source Data file.

Soft-clustering analysis classified rhythmic transcripts into four discrete clusters (Fig. 2b). Of the total rhythmic transcripts, 95% were clustered, with 1288, 431, 1143, and 1388 sequences assigned to clusters 1, 2, 3, and 4, respectively (Supplementary Data 3). Most rhythmic transcripts in clusters 1 and 4 showed acrophases during the day at ∼4-5zt and ∼6-8zt. Cluster 2, containing the least number of transcripts (10% of clustered rhythmic transcripts), was represented by sequences with acrophases occurring during the late night-early morning transition (∼21-1zt). Cluster 3 was mainly represented by transcripts with acrophases during the night at ∼16-20zt.

Pathway enrichment analysis showed that cluster 1 was enriched in the pathways for the spliceosome, DNA replication, and basal transcription factors (Fig. 2c, Supplementary Data 3), indicating the active regulation of mRNA and protein production during the day. Cluster 2 did not show significant pathway enrichment but exhibited significant gene ontology (GO) terms related to small molecule, lipid, and carbohydrate metabolic processes (Supplementary Fig. 1). Cluster 3 displayed enrichment in pathways associated with lysosome and various biochemical processes involved in starch, sugar, nitrogen, and amino acid metabolism, indicating higher night-time carbohydrate and amino acid metabolic activities in aphids. Cluster 4 was notably enriched in pathways related to purine metabolism, thiamine metabolism, and multiple pathways associated with human neurodegenerative diseases.

### Silencing diurnally rhythmic osmoregulatory genes impacts aphid performance

Osmoregulation is an important physiological activity to tolerate sugar-rich phloem sap during aphid feeding. Three osmoregulatory genes, *aquaporin 1* (AQP1) ^31^, *gut sucrase 1* (SUC1) ^30,38,39^, and *gut sugar transporter 4* ^40^, were previously reported to display diurnal patterns in *R. padi* ^16^. In our study, all three genes were confirmed to be diurnally rhythmic, with acrophases occurring at 15-22zt (Fig 2d-e, Supplementary Table 3). To determine whether the osmoregulatory genes contributed to diurnal variations in honeydew excretion, AQP1, and SUC1 were silenced by injecting aphids with dsRNA targeting each gene. A significant silencing of AQP1 and SUC1 was confirmed 24 hours post-injection (hpi) (Supplementary Fig. 2a-b) with a knockdown efficiency of 44% and 50%, respectively. Silencing AQP1 and SUC1 increased aphid mortality over time, and reduced nymph production by 31% and 66% at 96 hpi, respectively (Fig. 2f-g). Moreover, silencing the two genes markedly reduced honeydew excretion and abolished rhythms in the number of honeydew droplets produced by aphids between the day and night (Fig. 2h). This could suggest that the diurnal variation in these two genes is responsible for the diurnal variation of honeydew excretion and osmoregulation seen in *R. padi*.

### Diurnal rhythmicity in *R. padi* salivary effectors

Given the observed diurnal rhythm in the E1 salivation phase, we sought to determine the presence of diurnal rhythms in the expression of putative aphid salivary effectors. Previously, 156 genes were identified as putative salivary effectors in *R. padi* ^23^. Our study retrieved 93% of putative effectors (145 genes represented by 264 transcripts) by searching all transcripts against the *R. padi* putative effector dataset (Supplementary Data 4). Among them, 42 exhibited diurnal rhythmicity, with 28 exhibiting acrophases at 6-8zt during the day and 14 with acrophases at 18-19zt and 22-23zt at night (Fig. 3a, Supplementary Table 4).

**Figure 3.**
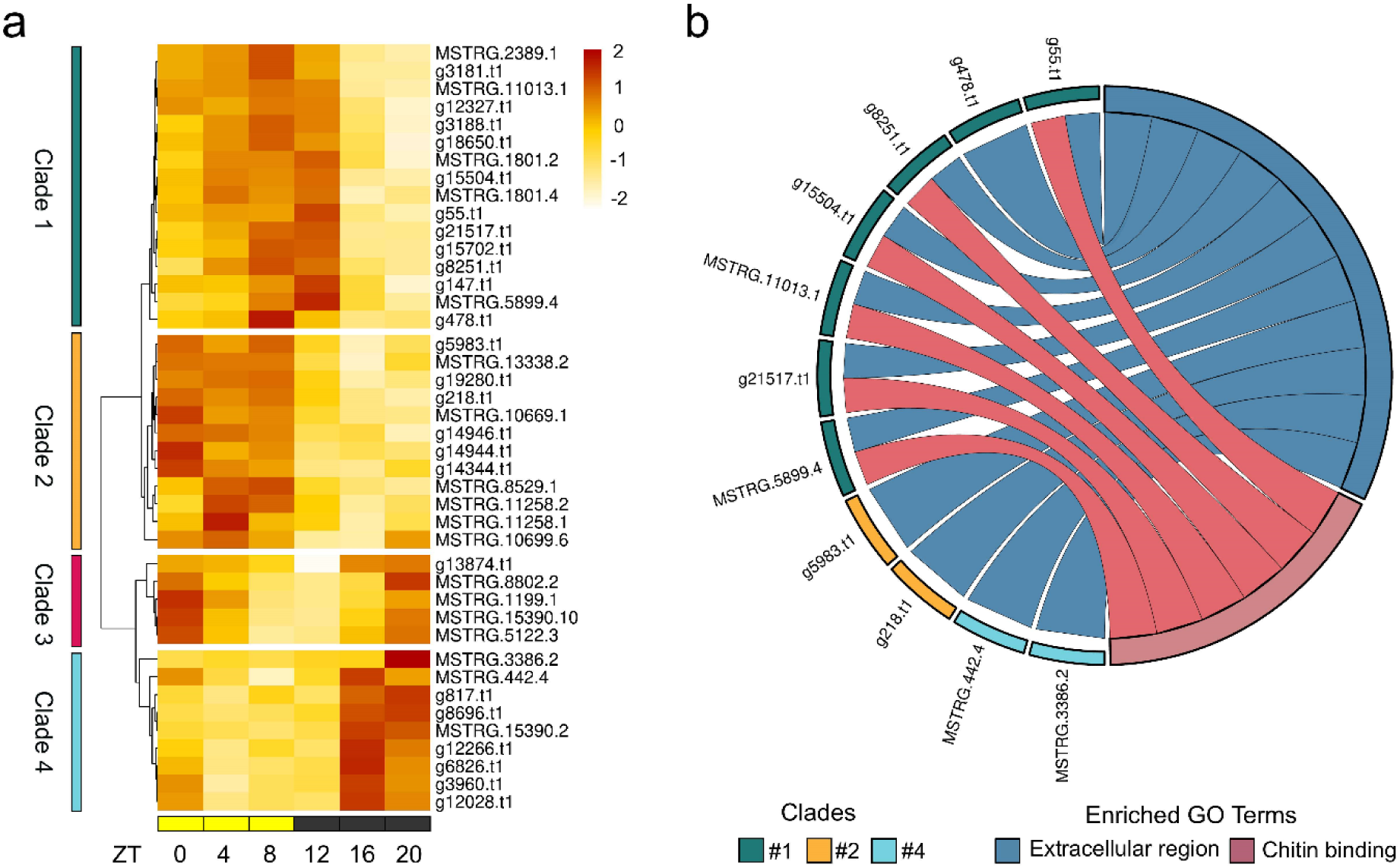
Diurnally rhythmic putative salivary effectors in *R. padi*. **a** Heatmap showing temporal gene expression patterns of 42 diurnally rhythmic putative salivary effectors. Each row and column represents the salivary effectors and zeitgeber time (ZT). The hierarchical clustering of gene expression data was generated using one minus Spearman correlation metric based on the average linkage method. Color scale represents the standardized expression values (Z-score) across samples. **b** Enriched gene ontology (GO) terms associated with diurnally rhythmic putative salivary effectors. Node color corresponds to the clade number in the heatmap (**a**). Source data and full statistical summary are provided in the Source Data file.

The expression levels of these rhythmic putative effectors were further examined in *R. padi* heads compared to the bodies (body samples excluded tissue from heads and unborn nymphs). We confirmed that the expressions of all rhythmic putative effectors were significantly higher in the heads, except for the putative effector MSTRG.15390.2, which lacked assigned count values across head and body samples (Supplementary Fig. 3). Hierarchical clustering of the rhythmic putative effectors identified four main clades, each showing distinct times of peak expression (Fig. 3a). Transcripts in clades 1 and 2 peaked in expression at the transition from day to night and the early portion of mid-day, respectively. In contrast, transcripts within clade 3 exhibited peak expressions at the end of the night and early morning, while clade 4 consisted of transcripts with expression peaking towards the end of the night. Notably, 43% of rhythmic putative effectors with peak expressions occurring at 12-16zt in clade 1 and 4 coincided with the timing of increased E1 salivation, including a well-characterized salivary effector - C002 (AphidBase accession - g12266.t1).

GO enrichment analysis highlighted two significantly enriched terms for all putative effectors: extracellular region and chitin-binding (Fig. 3b). The former supports a secretory role for these putative effectors, and the latter implies their potential time-dependent functions in evading or interfering with chitin-triggered plant immunity ^41,42^. These findings provide evidence of diurnal rhythms in salivary effector expression, with a subset peaking in expression at the same time as increased nighttime salivation. It also suggests that different blends of salivary effectors are secreted into the plants at different times of the day, providing novel evidence of diurnal rhythmicity in salivary effector gene expression in aphid species.

### Key salivary effectors identified for functional analysis

To elucidate the functions of diurnally rhythmic salivary effectors in aphid-plant interactions, we selected C002 and a novel putative effector, E8696 (AphidBase accession - g8696.t1), based on their high amplitude values (i.e. half the difference between the highest and lowest points of a rhythmic cycle), positioning them among the top five compared to other rhythmic putative effectors (Fig. 4a, Supplementary Table 4). According to our soft-clustering analysis, C002 and E8696 were assigned to cluster 3 with membership scores of 0.895 and 0.979, respectively (Supplementary Data 3). Their high membership scores designate them as core members within the cluster ^43^. Additionally, both genes also showed significantly higher expression in *R. padi* heads compared to bodies (Fig. 4b) and peaked in expression at 16zt (midnight) (Fig. 3c-d).

**Figure 4.**
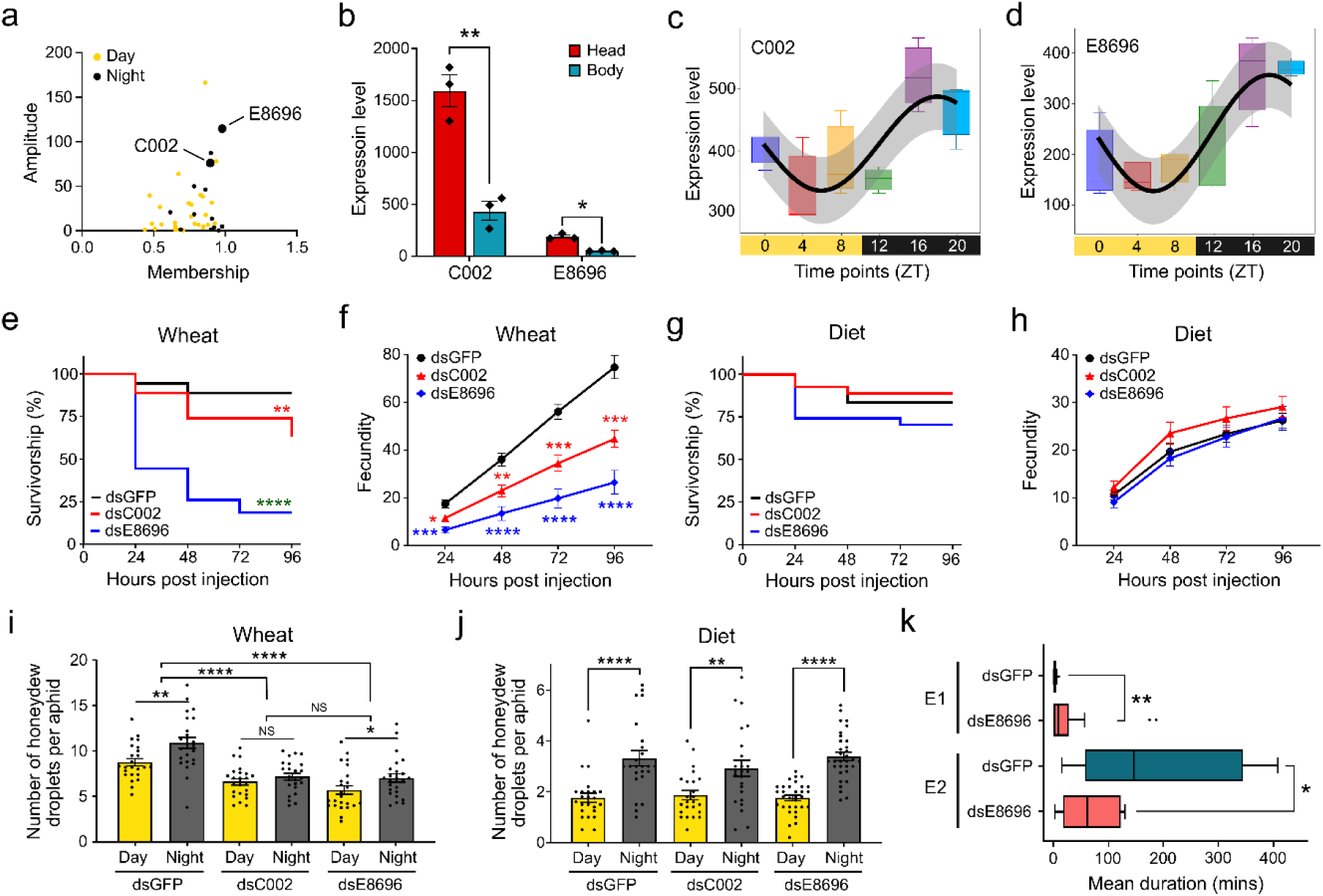
Aphid performance on wheat and artificial diet after silencing diurnally rhythmic putative salivary effectors. **a** Identification of key putative salivary effectors for functional analysis. Yellow and black dots represent diurnally rhythmic effectors with acrophases during the day and night. **b** Expression levels of putative salivary effector genes, C002 and E8696, in *R. padi* heads (n=3) and bodies (n=3, without heads and nymphs) are compared using a two-tailed Welch’s *t*-test (C002: t=6.532, df=3.218, \**p=*0.0059 and E8696: t=8.528, df=2.072, \*\**p*<0.0121). **c, d** Expression profiles of C002 (**c**) and E8696 (**d**) over a 24h period (n=3 per time point). Solid lines, boundaries, and whiskers of each box indicate the median, 25th, and 75^th^ percentiles, and 1.5 interquartile ranges, respectively. Diurnal rhythmicity of C002 (*p*=0.008) and E8696 (*p*=0.001) was determined by Cosinor model. The curve line and shaded area represent the fitted periodic sinusoidal curve and 95% confidence band, respectively. **e** Kaplan-Meier survival curves and **f** fecundities of aphids on wheat plants after treatment with dsRNA C002 (dsC002, n=3/9, representing the number of replicates for survival and fecundity, respectively), dsRNA E8696 (dsE8696, n=3/9), and dsRNA GFP (dsGFP, n=6/12, negative control). **g** Kaplan-Meier survival curves and **h** fecundities of aphids on diets after treatment with dsC002 (n=3/9), dsE8696 (n=3/11), and dsGFP (n=6/12). Each replicate (n) contains five individual aphids. Significant differences in survivorship and fecundity were tested using a log-rank test with Benjamini-Hochberg adjustment and a repeated measures ANOVA with Dunn’s multiple comparisons test, respectively (\**p*<0.05, \*\**p*<0.01, \*\*\**p*<0.001, \*\*\*\**p*<0.0001). **i, j** The number of honeydew droplets produced per aphid after treatment with dsC002, dsE8696, and dsGFP on wheat plant (**i**, n=6 per treatment group) and diet (**j**, dsC002: n=6, dsE8696: n=8, and dsGFP: n=6). Each replicate (n) contains five individual aphids. Yellow and black bars represent the mean (±SE) number of honeydew droplets produced at the end of each 12h day and night cycle for 4 days (Mann-Whitney test, \**p*<0.05, \*\**p*<0.01, \*\*\*\**p*<0.0001). **k** Aphid feeding activity after treatment with dsE8696 (n=13) and dsGFP (n=15). Solid lines, boundaries, and whiskers indicate the median, 25^th^, and 75^th^ percentiles and 1.5 interquartile ranges, respectively. Points indicate outliers. Asterisk(s) denote significant differences (Mann-Whitney test, E1: \*\**p=*0.0071 and E2: \**p=*0.0195). Source data and full statistical summary are provided in the Source Data file.

Sequence analysis revealed that E8696 encodes for a protein of unknown function with a predicted signal peptide (probability score: 0.953, cleavage site: amino acids 24-25) (Supplementary Fig. 4a). Phylogenetic analysis demonstrated the high conservation of this novel effector in aphid species, particularly in close relation to another cereal aphid, the corn leaf aphid, *R. maidis* (Supplementary Fig. 4b).

### Disrupting diurnally rhythmic salivary effectors alters feeding dynamics and performance

We further determined the role of C002 and E8696 in aphid feeding and performance through RNA interference. A significant knockdown of C002 and E8696 expression was confirmed at 24 hpi, with significant C002 silencing sustaining until 48 hpi (Fig. S2c-d). Silencing efficiencies at 24 and 48 hpi were 37% and 27% for C002 and 29% and 26% for E8696, respectively. Silencing C002 reduced aphid survivorship and fecundity on wheat (Fig. 4e-f), consistent with previous findings ^24,26,44^. Specifically, C002 silencing caused 29% aphid mortality and a 56% reduction in fecundity at 96 hpi compared to dsGFP control. Similarly, silencing of E8696 reduced aphid survivorship and fecundity, causing a 79% mortality and 86% reduction in fecundity at 96 hpi. To uncouple the influence of plant-derived cues on aphid feeding, we also assessed aphid performance on artificial diets after silencing the two effector genes. In contrast to the results observed on wheat plants, silencing either gene did not affect aphid survivorship and fecundity (Fig. 4g-h).

On plants, silencing either effector gene reduced overall honeydew excretion, with only C002 silencing disrupting the diurnal rhythm in honeydew excretion (Fig. 4i), suggesting its time-oriented roles in aphid feeding on plants. Nighttime honeydew excretion decreased by 1.7- and 1.4-fold compared to the day after silencing C002 (4.1 vs 2.4) and E8696 (4.1 vs 2.9). In contrast, silencing had no impact on honeydew excretion on artificial diets, and aphids consistently generated more honeydew droplets at night than during the day (Fig. 4j). Furthermore, we found no diurnal variation in nymph production post-silencing on both plant and diet (Supplementary Fig. 5). However, the overall number of nymphs was markedly lower compared to dsGFP controls on the plant, but not on the diet.

Disturbed aphid feeding activities were also confirmed using EPG after silencing E8696. Aphids exhibited an increase in the average duration (total duration/number of each event) of the salivation phase (E1), accompanied by a concurrent decrease in the phloem ingestion phase (E2) (Fig. 4k, Supplementary Data 5). These findings confirm the vital role of E8696 in aphid feeding behavior and performance on host plants.

## Discussion

The cyclical patterns of environmental cues, dictated by the earth’s light-dark cycle, significantly influence the physiology and behavior of organisms, including crop plants and their pests ^1,6^. While studies have begun to show the prevalence of diurnal rhythms among aphid pests, critical knowledge gaps persist regarding how these rhythms facilitate interactions between aphids and their hosts. Our study aimed to fill these knowledge gaps by exploring diurnal rhythms in behavior and gene expression in the bird-cherry oat aphid (*R. padi*), a major pest on cereals worldwide. We further characterized the impact of diurnally rhythmic osmoregulatory and salivary effector genes on *R. padi* behavior and performance.

Here, we found that aphids exhibited diurnal rhythms in feeding behaviors, particularly in honeydew excretion and the duration of phloem salivation (E1 phase). Contrary to previously documented patterns ^10,11^, *R. padi* excreted more honeydew droplets on wheat plants during the nighttime. This divergence in honeydew excretion patterns implies species-specific adaptations of aphids to host plants, possibly indicating differences in aphid osmoregulatory rhythms in response to phloem quality across host species. Notably, the diurnal honeydew excretion patterns persisted in *R. padi* on artificial diets, devoid of plant-derived cues, suggesting a potential internal circadian clock-driven mechanism. However, we cannot fully rule out the influence of plant-derived factors. Diurnal fluctuations in phloem sap composition, such as changes in sugar concentrations that affect osmotic potential or nutrient availability, could still modulate aphid osmoregulatory responses. Future studies experimentally manipulating both plant-derived cues and aphid physiological pathways will be critical to disentangle the relative contributions of internal and external drivers. It is important to note that our experiments were conducted in the absence of plant cues only. To determine the circadian clock’s involvement in regulating the rhythms identified in our study, all cues, including plant, light, and temperature, would need to be removed.

Remarkably, the pronounced nighttime increase in phloem salivation suggests that aphids may be responding to changes in plant defenses or phloem quality, both of which are known to exhibit diurnal variation in numerous plant species ^45–47^. We also found that the duration of phloem and xylem sap ingestion (E2 and G phases, respectively) showed no significant differences over a 24 h cycle. The desire for aphids to quickly reach and remain in the E2 phase regardless of timepoint, coupled with the strong diurnal variation in honeydew excretion, might suggest that aphids do not modulate feeding behavior as a tool for osmoregulation but rather rely on metabolic mechanisms to process and excrete excess sugar. The low occurrence and the lack of diurnal variation in the G phase further suggest that increased metabolism of ingested phloem sap, resulting in higher honeydew excretion, constitutes the primary mechanism for aphid osmoregulatory function over a 24 h period, with xylem ingestion only occurring in aphids that are severely dehydrated. These findings align with aphids’ nutritional and reproductive strategies, underscoring the importance of acquiring sufficient sugars and amino acids for population growth and survival.

Our time-course profiling of aphid transcriptome revealed that approximately 9% of the total assembled transcripts displayed diurnal rhythms. Most rhythmic transcripts showed acrophases 6 h after the start and end of the light cue (noon or 4zt and midnight or 16zt, respectively). The bimodal pattern of these acrophases suggests that noon and midnight are periods of high activity in aphids, with multiple physiological and behavioral processes occurring at or around these times. Similar diurnal rhythmicity in gene expression was observed in Drosophila and the malaria mosquito (*Anopheles gambiae*), estimating rhythmic expression in 10% and 15.8% of genes linked to various biological functions ^45–48^, respectively. Clustered rhythmic transcripts, showing peak expressions at specific times of the day, point to a transcriptional control mechanism, possibly by the circadian clock. Our enrichment analysis highlights the functional diversity within the different rhythmic gene clusters, particularly in metabolic activities between the day and night. Notably, the night-peaking clusters 2 and 3 were enriched in pathways related to the metabolism of carbohydrates, amino acids, lipids, and other small molecules. The upregulation of these metabolic pathways at night corresponds with the observed diurnal rhythms in honeydew excretion, suggesting a correlation between these metabolic processes and the regulation of honeydew excretion.

Previous studies show that silencing AQP1 and SUC1 in *M. persicae* led to elevated hemolymph osmotic pressure, accompanied by a decrease in aphid weight and fecundity ^32^. In our study, a parallel decline in aphid performance was observed: silencing AQP1 and SUC1 in *R. padi* resulted in increased mortality and reduced fecundity, highlighting their broad physiological significance for aphid homeostasis and life-history traits. Notably, both genes exhibit pronounced diurnal rhythms, suggesting peak osmoregulatory activity at specific times of day, potentially linked to changes in phloem sap composition. An important aspect of osmoregulation in aphids is the excretion of excess sugar in the form of oligosaccharides in honeydew ^27^. The overall reduction in quantity and the abolition of rhythms in honeydew excretion upon the silencing of AQP1 and SUC1 confirm their pivotal roles in shaping diurnal behavior of aphid honeydew excretion. While acknowledging their well-established broader physiological roles in maintaining osmotic balance and overall fitness, our findings strongly indicate that AQP1 and SUC1 are also key players in the diurnal regulation of aphid feeding and excretory processes.

While we found that *R. padi* spent similar amounts of time feeding from the phloem cells during both day and night, a significant increase in salivation was observed at the beginning of the night. This suggests that time-of-day variations in the host plant may prompt aphids to modulate host responses through salivation and the introduction of specific effectors. In this study, we use the term putative salivary effector to describe aphid-expressed, head-enriched, secreted-protein candidates whose secretion into host tissue and host targets have not yet been experimentally validated. Indeed, we found clear evidence showing diurnal rhythms in the expression of several putative salivary effector genes, including C002. The dynamic expression patterns of these rhythmic putative effectors suggest that different blends of effectors may be introduced into the plant for modulating time-of-day specific host plant responses. However, rhythmic transcript accumulation may not necessarily indicate proportional changes in protein abundance or secretion into saliva and host tissues. Future studies integrating protein-level detection and localization approaches will be required to determine whether rhythmic gene expression corresponds to rhythmic protein secretion.

To further confirm the time-dependent role of these rhythmic putative effectors in aphid-plant interactions, we silenced the C002 and E8696 effector genes, both showing peak expression at night. Silencing these two genes resulted in high mortality and reduced fecundity in aphids when feeding on host plants. However, no such trend was observed when the same experiment was performed on artificial diets, underscoring the crucial role of these putative effectors in facilitating aphid colonization of host plants. Strikingly, silencing these two effectors also resulted in a more pronounced reduction in nighttime honeydew excretion compared to the day on host plants. However, it had no impact on diurnal honeydew excretion on artificial diets. This affirms their significant roles in optimizing aphid feeding strategies and potentially enhancing their adaptation to the host plant’s physiological changes during the night. Future research is warranted to elucidate the molecular targets and mechanisms underlying these phenotypes and whether the rhythmic activity of these putative effectors is driven by the aphid’s circadian clock, phloem sap compositions, plant defenses, environmental changes across the diurnal period, or some combination of these factors.

In summary, our findings reveal pronounced diurnal variation in the behavior and physiology of a major agricultural pest, underscoring the importance of time as an organizing factor in insect– plant interactions. Considering daily rhythms in aphid feeding, metabolism, and physiological performance provides a framework for moving beyond static, time-averaged views of pest biology. Temporal mismatches between aphid activity and host plant physiological states, including circadian regulation of plant defenses, may create exploitable windows of vulnerability that could be leveraged through breeding, cultural practices, or integrated management strategies. In addition, diurnal variation in aphid osmoregulation and stress tolerance suggests that interventions targeting critical physiological processes during specific phases of the day–night cycle may reduce fitness and population growth while limiting continuous selection pressure. By strategically leveraging the daily rhythms that mark periods of strength and vulnerability in both crop plants and their pests, we envision entering an era of “chronoculture,” ^6^ in which harnessing circadian rhythms contributes significantly to enhancing crop yield and promoting sustainability.

## Methods

### Aphid colonies and plant growth conditions

A colony of bird cherry-oat aphids (*Rhopalosiphum padi*) was initiated with aphids collected from greenhouses at the Plant Growth Facility at Colorado State University, Fort Collins, and has been maintained in the laboratory since 2019. *R. padi* was reared on three-week-old barley seedlings (*Hordeum vulgare* var. AC Metcalfe). Colony plants were replenished every two to three weeks or when overcrowding occurred. All plants were grown in PRO-MIX BX General Purpose Growing Medium (Premier Tech Horticulture, Quakertown, PA) in mylar plant growth tents at 60-70% relative humidity, temperature of 24 ± 1°C and maintained under a short-day photoperiod of 12L:12D at photosynthetically active radiation (PAR) of 460 μmol/m^2^/s. Plants were watered three times per week and fertilized with Miracle Gro® (Scott’s Co. LLC, Marysville, OH, USA) once a week. All experiments were performed on wheat (*Triticum aestivum* var. Chinese Spring) plants that were 14-21 days old or at the two-leaf stage (Zadoks stage, Z1.2) ^48^.

### Aphid age synchronization

All experiments used age-synchronized aphids to limit any age-related effects on aphid behaviors or gene expression ^49^. Briefly, adult aphids from the colony were placed into a feeding chamber containing the second and third leaves of a two-week-old wheat plant. The feeding chamber was made from a 50-ml falcon tube with the bottom side cut open and the lid modified for ventilation using aphid-proof mesh. Leaves were inserted into the tube from the bottom opening and sealed with a cotton ball. After 48 h, adult aphids were removed, and nymphs were allowed to develop to adulthood on the plant for five days.

### Aphid feeding assay on wheat and artificial diet

Five age-synchronized aphids were placed into a 55 mm Petri dish (V.W.R., Radnor, PA) modified to contain leaf 2 of a wheat plant (Z1.2) (Fig. 5a). Nymph count and honeydew droplet production were recorded at the onset of day and night over 48 h, adhering to a 12L:12D photoperiod. Discrete honeydew droplets were counted instead of using a staining and area calculation method. This choice was made because droplets exhibit uniform size and were easily visible to the naked eye. However, their translucent color made capturing them in photographs and measuring the area challenging. After each measurement, nymphs were carefully removed, and the honeydew-soiled lid was replaced with a fresh one. Concurrently, feeding assays were conducted using an artificial diet designed for *R. padi* ^50^. An artificial feeding chamber was constructed under sterile conditions and consisted of 58 mm Petri-dish lids with Parafilm sachets containing 1.5 ml of diet. Five age-synchronized aphids were placed into the artificial feeding chamber (Fig. 5a) and the number of nymphs and honeydew droplets were measured as described above. Both experiments were independently conducted on two occasions, with five replicates on each occasion (n = 10 for each experiment on wheat or diet). Statistical differences between treatments were tested using the Mann-Whitney test (*P* < 0.05) in Graphpad Prism v10.

**Figure 5.**
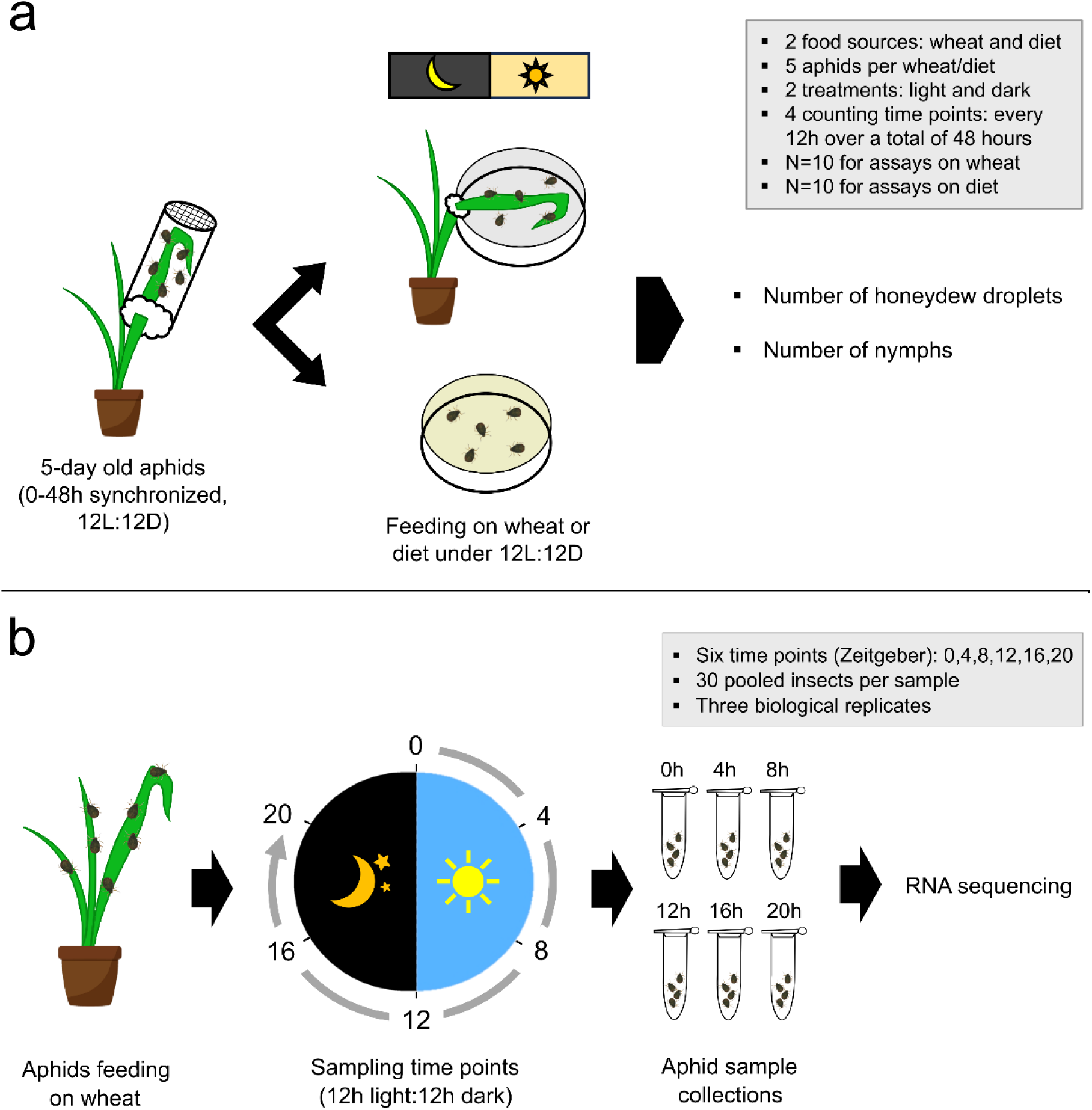
**a. Diurnal behavioral activities of *R. padi* on wheat plant and artificial diet.** Five age-synchronized adult females were placed on a wheat leaf or diet under 12h:12h, light:dark cycles. The number of honeydew drops and nymphs produced by aphids was recorded at the end of each night and day for two days. Each experiment on wheat or diet contains a total of 10 biological replicates. **b. Diurnal time-course experiment.** Aphid samples were collected every four hours over a 24h day-night cycle (n = 3 per timepoint).

### Electrical penetration graph

Adult *R. padi* feeding behavior was recorded on wheat seedlings (Z1.2) using the electrical penetration graph (EPG) ^51^ technique on a GIGA 8 complete system (EPG systems, Wageningen, Netherlands). The recordings were performed every 4 h over a 24 h period starting at 0zt (a total of six time periods). Age-synchronized adult *R. padi* were starved for 1 h before wiring. Next, plant electrodes were inserted into the soil, and insect probes were adjusted to ensure contact between the plant surface and the insect. Aphids were only allowed to feed from the leaf 2, and only one aphid was recorded per plant. For recordings that were initiated in the night or scotophase (0zt, 16zt, and 20zt), all setup was performed with a red LED headlight (Energizer). The 0zt timepoint was set up in dark conditions while feeding behavior was recorded in light. The 12zt timepoint was set up in light conditions, with feeding behavior recorded in darkness. Photoperiod and the temperatures during the day and night were the same as those used for plant and aphid growing conditions. Both plants and aphids were discarded after each 4 h recording. EPG waveforms were analyzed using the Stylet+ software (v01.25, EPG systems, Wageningen, the Netherlands). The analysis of each feeding experiment aimed to determine the time allocation across four primary phases: the pathway or probing phase (C), non-probing phase (NP), salivation phase (E1), phloem sap ingestion phase (E2), and xylem phase (G). Additionally, many EPG parameters that characterize aphid probing and ingestion ^52^ were determined to develop a comprehensive picture of aphid feeding over a 24 h period. EPG parameters were calculated using the discoEPG package integrated into the web-based R/shiny app (https://nalamlab.shinyapps.io/test/). The source code used for discoEPG software development is available on GitHub platform (https://github.com/nalamvj/EPG_ANALYSIS). There were 25 replicates for 0zt, 21 replicates for 4zt, 26 replicates for 8zt, 23 replicates for 12zt, 29 replicates for 16zt, and 16 replicates for 20zt. Recordings without feeding or probing events and those with more than 70% of time spent in NP + derailed stylet phase (F) + G were excluded from the analysis. Statistical differences between the six time periods of each EPG phase were tested using the Kruskal–Wallis test with Bonferroni adjustment for *p*-values (adjusted *P* < 0.05). The rhythmicity of EPG phases was determined with the average total duration (minutes) of each recorded waveform across the different periods using a Cosinor-based oscillation detection method in DiscoRhythm v1.2.1 ^53^. EPG phases were considered significant rhythmic if their *p*-values were less than 0.05, and their corresponding oscillation parameters were estimated.

### Sample collection, RNA isolation, library preparation, and RNA Sequencing

A cohort of adult *R. padi* (∼100) were evenly distributed on wheat plants (Z1.2). The adults were removed after 24 h, and the nymphs were allowed to develop to adulthood for six days. These age-synchronized *R. padi* adults were collected from separate wheat plants every four hours over a 24 h period (12L:12D), starting at 0zt (when the lights were turned on) and ending at 20zt (Fig. 5b). A pool of 30 apterous aphids was collected per sample and extracted for total RNA using the Quick-RNA Miniprep Kit (Zymo Research, Irvine, CA) according to the manufacturer’s instructions. This time-course RNA-Seq experiment comprised three biologically independent replicates of aphid samples per time point. RNA quality and concentration were measured using a Nanodrop spectrophotometer (Thermo Scientific, Waltham, MA) and Agilent 2100 Bioanalyzer (Santa Clara, CA). Aphid RNA samples were sent to Novogene Corporation Inc. (Sacramento, CA) for cDNA library construction and RNA sequencing. The poly(A) mRNA-enriched libraries were constructed using NEBNext® Ultra™ II RNA Library Prep Kit (New England BioLabs, Ipswich, MA). Paired-end sequencing (2×150 bp) was conducted with the Illumina HiSeq 2000 platform. The raw sequencing reads have been deposited in the NCBI SRA database under the BioProject accession PRJNA979927.

### Genome-guided transcriptome assembly and read mapping

Raw sequence reads were trimmed for adapters and quality using fastp v0.23.3^54^. The reads with a Phred score less than 25 and lengths shorter than 100 bp were discarded. The high-quality reads from each library were mapped against the *R. padi* reference genome (v2.0) available on AphidBase (https://bipaa.genouest.org/is/aphidbase/) using Hisat2 v2.2.1. The mapped reads were assembled into transcripts using Stringtie v2.2.1. The assemblies from all 18 libraries were further merged to generate a reference transcriptome using the Stringtie -merge function ^55^. Reads representing each sampling timepoint were mapped back to this reference transcriptome to quantify the gene expression using Salmon v1.9.0 ^56^.

### Identification and clustering of diurnal rhythmic transcripts

The non-random rhythmicity of *R. padi* genes was determined with a Cosinor-based oscillation detection method in DiscoRhythm v1.2.1 ^53^. The temporal expression data of all transcripts identified in the whole-body *R. padi* (48,690) were utilized for detecting rhythmic signals. The expression data was filtered for constant or missing values (>25%) across the samples, resulting in 38,871 transcripts. Rhythm detection and estimation of oscillation parameters were performed as previously outlined. Soft clustering of all rhythmic sequences was carried out to identify major temporal patterns of rhythmic expression using the R package, mFuzz v2.56.0 ^43^. Membership values ranging from 0 to 1 were assigned to clustered transcripts. Sequences with a membership value > 0.7 were considered core members of each cluster ^57^.

### Gene ontology and pathway enrichment analyses

All clustered sequences of rhythmic transcripts in *R. padi* were blastx-searched against the NCBI non-redundant (nr) protein database using OmicsBox v2.2 with an E-value cut-off of 1e-5 ^58^. Sequences with assigned gene ontology (GO) terms were categorized into biological process, molecular function, and cellular component categories. Further, these clustered sequences were annotated for the Kyoto Encyclopedia of Genes and Genomes (KEGG) orthologs and linked to corresponding KEGG reference pathways. GO and KEGG pathway enrichment analyses were performed for rhythmic transcripts within each of the four gene clusters and the rhythmic putative salivary effector transcripts using a two-tailed Fisher’s exact test with an FDR cut-off of 0.05 in OmicsBox ^58^. The entire *R. padi* transcriptome was used as the background gene set for both enrichment analyses.

### Identification of diurnally rhythmic osmoregulatory and putative salivary effector genes

Osmoregulatory genes in *R. padi* were identified as described previously ^16^. Briefly, homologs for these genes were identified by querying *Acyrthosiphon pisum* gene sequences with the *R. padi* reference genome (v2.0) on AphidBase ^59^ using Blastp. Further validation involved searching protein sequences with DeepTMHMM ^60^ and SMART ^61^ to determine the presence of sequence features. All *R. padi* transcripts were blastx-searched against the curated putative salivary effector nucleotide database for *R. padi* ^23^ using Blast+ v2.13.0 with an E-value cut-off of 1e-5. Stringent criteria were applied, with thresholds set at 80% for identity and 50% for coverage to identify putative salivary effectors. To quantify tissue-specific expression of these rhythmic putative effectors, RNA-Seq reads of *R. padi* heads and bodies (not including heads and nymphs) were retrieved through the European Nucleotide Archive under the project accession number of PRJEB9912 ^23^. Sample accessions for head and body datasets are as follows: SAMEA3505144, SAMEA3505145, SAMEA3505146, SAMEA3505147, SAMEA3505148, SAMEA3505149. Reads were trimmed for adapters and quality and mapped against the previously assembled *R. padi* reference transcriptome using the abovementioned pipeline. The diurnal rhythmicity of identified osmoregulatory and putative effector sequences was tested using the Cosinor model described above.

### Sequence and phylogenetic analyses of a novel putative salivary effector

The full-length coding sequence of E8696 was queried against the NCBI nr protein database with organism specified as aphids. There were 25 sequence hits in the database for E8696. The top hit of each aphid species was retrieved and used for sequence analysis (Supplementary Table 5). Signal peptide, transmembrane helix, and other conserved domains were analyzed using SignalP v6.0, DeepTMHMM, and NCBI Conserved Domain Search ^62–64^. Multiple sequence alignments were performed using MUSCLE, and the phylogenetic trees were built using the maximum likelihood method with GTR+G as the best-fit substitution model in MEGA 11 software ^65^. Node support rates were assessed with 1,000 bootstrap replicates.

### Design and synthesis of double-stranded RNAs

Sequence analysis was performed on the four target genes: AQP1, SUC1, C002, and E8696. The open reading frames and potential conserved domains were identified using the NCBI ORFfinder and Conserved Domain Search tools ^64^. The nucleotide sequence of each gene was evaluated to identify optimal target regions of double-stranded RNA (dsRNA). Corresponding primers were designed using the E-RNAi web tool ^66^ (Supplementary Table 6). The specificity of dsRNA primers was confirmed through a search against the NCBI *R. padi* nr database using the Primer-BLAST tool and in PCR, followed by agarose gel electrophoresis. For dsRNA synthesis, total RNA was extracted from a pool of ∼30 *R. padi* containing all five developmental stages using Direct-zol RNA Microprep Kit (Zymo Research, Irvine, CA) according to the manufacturer’s protocol. cDNA was synthesized from 1 µg of DNase-treated RNA using the Verso cDNA Synthesis Kit (Thermo Scientific, Waltham, MA). The dsRNA sequence region of each gene was amplified with the gene-specific primer pair using Q5 high-fidelity PCR (New England BioLabs, Ipswich, MA). The sense and antisense dsRNA sequence fragments conjugated with the T7 promoter sequence were amplified separately using the previous PCR amplicons as templates. All PCR products were confirmed for unique and accurate amplification in agarose gels and purified using GeneJET PCR Purification Kit (Thermo Scientific, Waltham, MA). dsRNA was synthesized with 1 µg of purified PCR products using the T7 RiboMAX Express RNAi System, according to the manufacturer’s instructions (Promega, Madison, WI). dsRNA targeting the green fluorescent protein (GFP) gene was synthesized following the protocol described above and used as a negative control. The quantity and integrity of dsRNA were measured by Nanodrop spectrophotometer (Thermo Scientific, Waltham, MA) and agarose gel electrophoresis.

### Microinjection of aphids with dsRNA

Injection needles (Drummond Scientific Co., Broomall, PA) were produced by pulling with a P-1000 micropipette puller (Sutter Instrument Co. Novato, CA) using optimized settings: heat=500, pull=100, velocity=40, delay=100, pressure=500, and ramp=493. The glass needle, filled with 5 μl of dsRNA, was assembled onto a Nanoject III microinjector (Drummond Scientific Co., Broomall, PA). Age-synchronized aphids were immobilized on a sticky tape strip (ventral side up) and injected with 180 ng of dsRNA (40 nl of 4.5 μg/μl) through the flexible membrane between the coxa of the hind leg and thorax. Immediately after injection, aphids were carefully detached from the tape using a wet paintbrush and transferred to a feeding chamber containing wheat leaves. The injection process was repeated until the desired number of aphids was achieved. The same procedure was applied to the other dsRNA treatment groups. The injected aphids were incubated at 25°C, 12L:12D photoperiod. Each experiment was independently replicated three times.

### Gene silencing in aphids

Five aphids, collected at 24- and 48-hour post-injection from three biological replicates, were subjected to RNA extraction using the Direct-zol® RNA Microprep Kit as described above. cDNA was synthesized from 500 ng of RNA and diluted 4-fold with nuclease-free water. Quantitative PCR (qPCR) was employed to measure the expression of the target gene and the housekeeping gene, Actin (AphidBase accession - g9633.t1). Gene-specific qPCR primers were designed using the NCBI Primer-Blast tool (Supplementary Table 6). A duplicate of 10 µl SYBR Green Supermix reaction (Bio-Rad, Hercules, CA) containing 2 μl of cDNA template was prepared and analyzed using CFX Connect Real-Time instrument (BioRad) with the following thermocycling conditions: 95°C for 30s, followed by 40 cycles of 95°C for 10s, 55°C for 30s, 72°C for 30s, with a final melt curve of 95°C for 30s, 65°C for 5s increasing to a final temperature of 95°C. The relative normalized expression of each target gene was estimated using the Pfaffl method ^67^. Statistical differences in gene expression between treatments at each time point were assessed using a two-tailed Welch’s *t*-test (*P* < 0.05) in Graphpad Prism v10.

### Aphid performance and feeding assays on wheat and artificial diet after gene knockdown

RNA interference (RNAi) was used to functionally characterize two osmoregulatory genes, aquaporin 1 (AQP1) and gut sucrase 1 (SUC1). Nine aphids injected with dsRNA AQP1 (dsAQP1) or dsRNA SUC1 (dsSUC1) were placed into a feeding chamber containing wheat leaves (Fig. 5a). Aphid mortality and fecundity (cumulative nymph count) were recorded every 24 h for 4 days. Aphids injected with dsRNA GFP (dsGFP) were the negative control. The number of honeydew droplets produced by five dsRNA-injected aphids on wheat plants were recorded at the beginning of the day and night for 4 days under a short-day photoperiod as described previously (Fig. 5a). Similar experimental setups were replicated for two target salivary effector genes, C002 and E8696, to measure aphid mortality and fecundity. Additionally, five aphids injected with dsRNA C002 (dsC002) or dsRNA E8696 (dsE8696) were placed on an artificial diet (Fig. 5b) and monitored for mortality and fecundity every 24 h for 4 days. To further assess daily changes in honeydew excretion and fecundity, five aphids after 24 h post-injection with dsC002 or dsE8696 were placed on a wheat leaf or artificial diet. The numbers of honeydew droplets and nymphs were recorded as previously described (Fig. 5a). All experiments were independently repeated three times. Kaplan-Meier survival curves were generated for dsRNA-injected aphids and compared for statistical differences by the Mantel-Cox log-rank test with Benjamini–Hochberg false discovery rate adjusted *p*-values (adjusted *P* < 0.05) using the Survminer v0.4.9. The statistical difference in the nymph count of dsRNA-injected aphids was determined using a repeated measures ANOVA test with Tukey’s multiple comparisons (adjusted *P* < 0.05). The normality of input data was confirmed using the Shapiro-Wilk test (*P* > 0.05). Statistical differences in honeydew and nymph productions between treatments were tested using the Mann-Whitney test (*P* < 0.05) and Kruskal-Wallis test, followed by Dunn’s multiple comparison tests (adjusted *P* < 0.05). The statistical analyses were performed using Graphpad Prism v10 and R v4.2.1.

### Electrical penetration graph after silencing E8696

Age-synchronized *R. padi* were injected with dsE8696 or dsGFP as described above. Twenty-four hours post microinjection, individual aphids were wired and subjected to EPG recording to document their feeding activities on wheat plants (Z1.2) as described previously. Recordings started at 0zt, and feeding behavior was recorded for 8 h for each aphid. The experiment included 13 replicates for dsE8696 and 15 replicates for the dsGFP control group. EPG parameters were calculated as previously described, and statistical differences between the two treatments were tested using the Mann-Whitney test (*P* < 0.05).

### Statistical analysis

The statistical tests employed for assessing significance between treatments were described in the respective sections above. All statistical analyses were performed in Graphpad Prism v10 and R v4.2.1.

## Data availability

The raw RNA sequencing reads of *Rhopalosiphum padi* have been deposited under the BioProject accession number PRJNA979927 in the Sequence Read Archive (SRA). Reference genome of *R. padi* (v2.0) can be found in AphidBase (https://bipaa.genouest.org/is/). All other data supporting the findings of this study are available in the article, Supplementary Information files, Source Data file, or from the corresponding author upon request. Source data are provided with this paper.

## Code availability

All code and software sources used in this study are described in the “Methods” section with corresponding citations of references, links to source codes, and Source Data file.

## Acknowledgments

We thank Joely Corrales for her assistance with planting and aphid synchronization. Drs. Jan Leach, Micky Eubanks, and Punya Nachappa for internal review of the manuscript. The project was funded by the United States – Israel Binational Science Foundation grant (Proposal#202181, PIs Vered Tzin and Vamsi Nalam), Research Capacity Fund (COL0044) from NIFA and start-up funds from Colorado State University to Vamsi Nalam.

## Author Contributions

J.H., D.K., V.T, and V.J.N designed the research; J.H., D.K., Y.G., and M.C. performed the research; J.H., D.K., M.C., Y.G., V.T, and V.J.N analyzed the data; and J.H., D.K., V.T, and V.J.N wrote the paper.

## Competing Interests

The authors declare no competing interests.

## Notes

### Competing Interest Statement

The authors have declared no competing interest.

### Summary of Updates

Terminology updated to indicate the E8696 as a putative salivary effector; Figure 4 legend revised; Discussion revised to indicate limitations of this study.

