## Supplementary information for "Diurnal rhythmicity in metabolism and salivary effector expression shapes aphid performance on host plants"

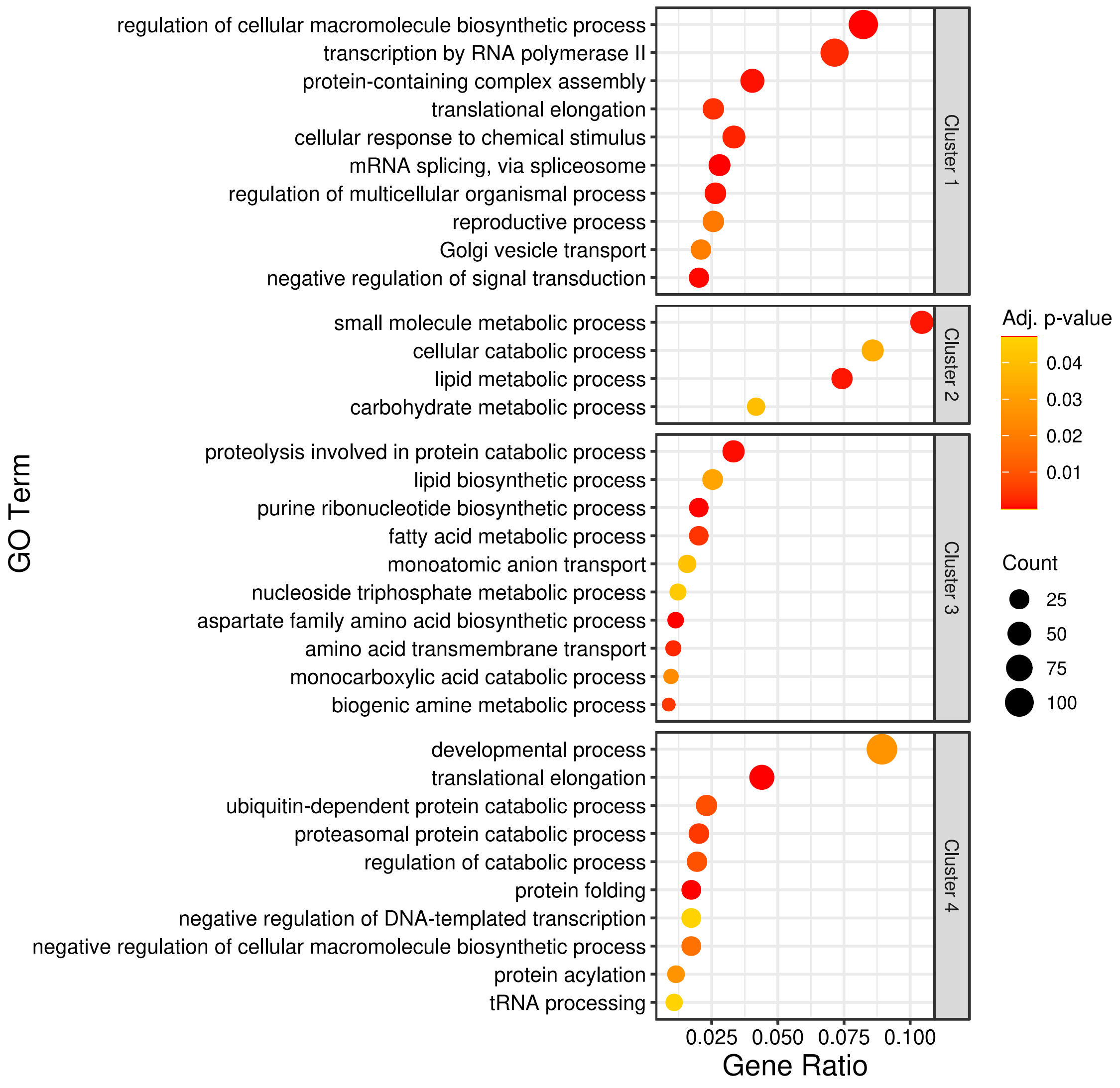


**Supplementary Figure 1.** Gene ontology (GO) enrichment analysis of diurnally rhythmic transcripts in each mFuzz cluster. The top-10 enriched GO terms in the category of biological process are displayed in the dot plot. Circle size and color represent the number of genes enriched in a GO term and Benjamini-Hochberg adjusted *p*-value, respectively. Gene ratio means the percentage of diurnally rhythmic transcripts of a cluster in the given GO term. Source data and full statistical summary are provided in the Source Data file.

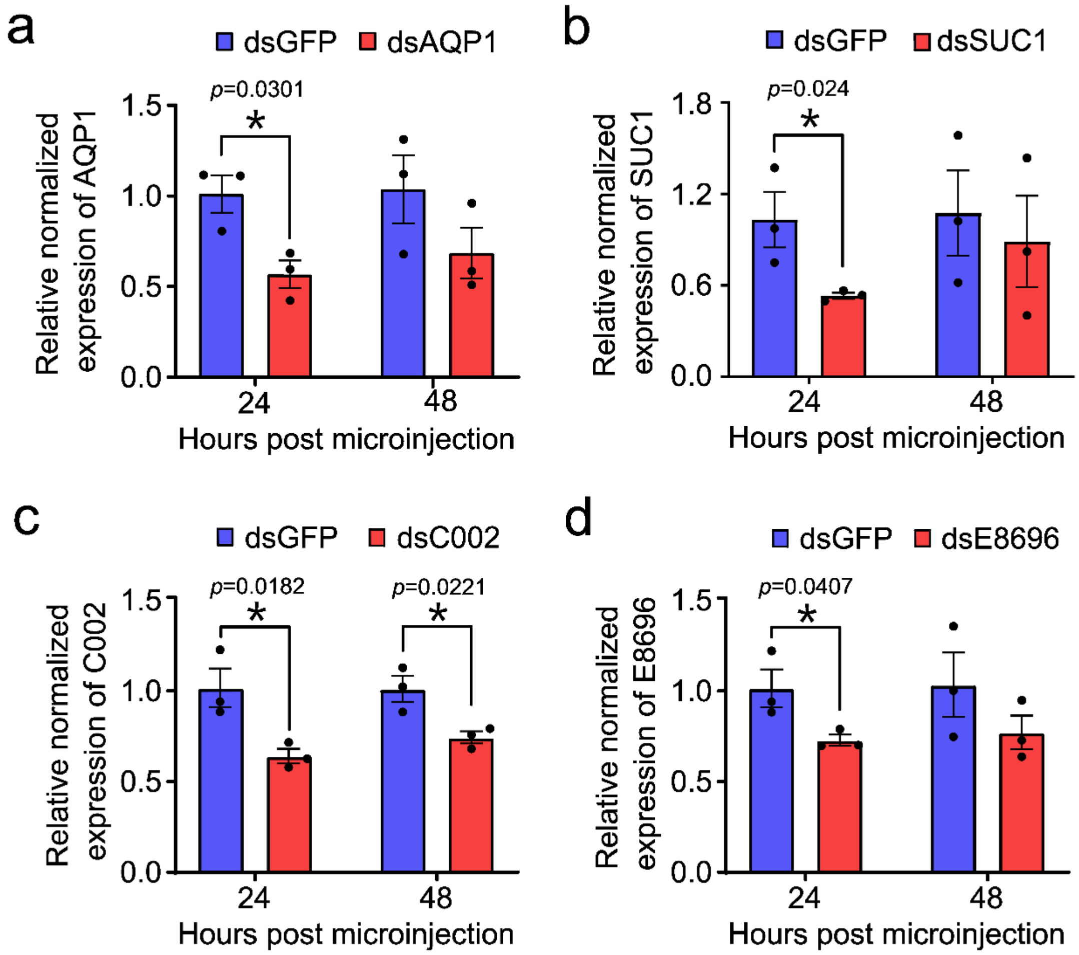


**Supplementary Figure 2.** Gene knockdown after injection with dsRNA. Knockdown efficiencies of (**a**) aquaporin 1 (AQP1), (**b**) gut sucrase 1 (SUC1), (**c**) C002, and (**d**) E8696 were determined after 24- and 48-hour post injection with dsAQP1 (n = 3 per timepoint), dsSUC1 (n = 3 per timepoint), dsC002 (n = 3 per timepoint), and dsE8696 (n = 3 per timepoint). Statistical difference in gene expression between treatments was tested using a two-tailed, Welch's *t*-test (*$p < 0.05$). Source data and full statistical summary are provided in the Source Data file.

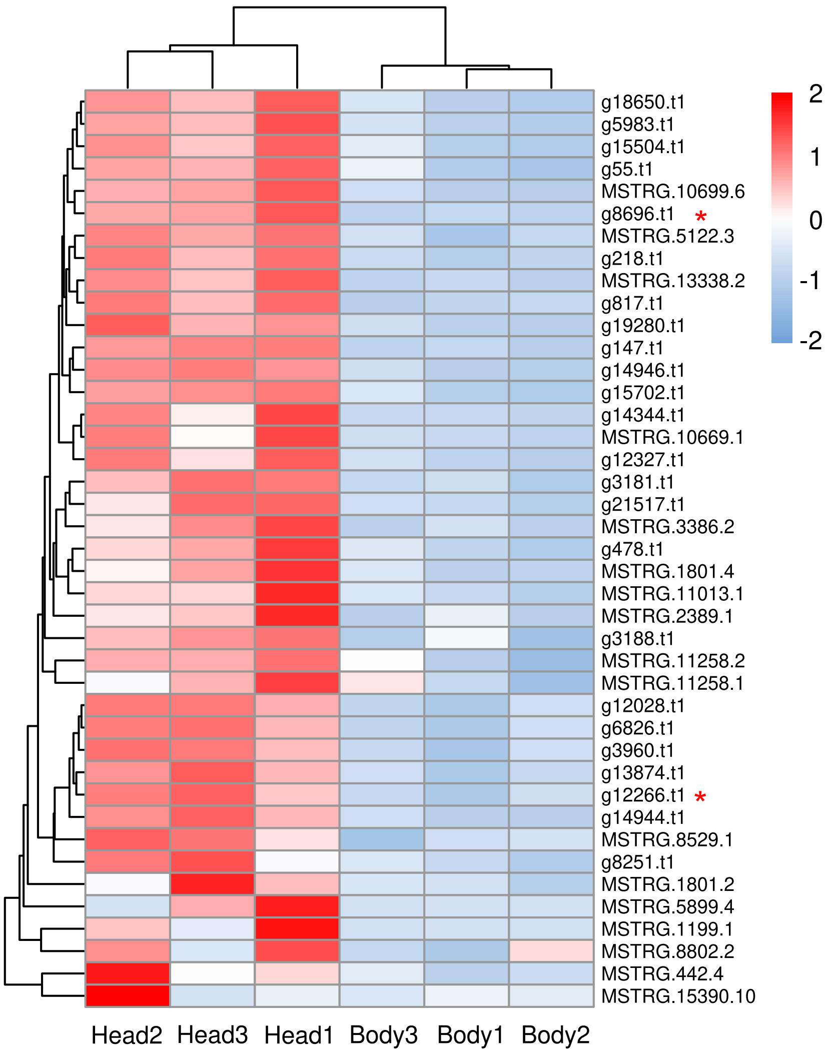


**Supplementary Figure 3.** Heatmap showing the expression levels of 42 diurnally rhythmic salivary effectors between *R. padi* heads and bodies. Raw RNA-Seq reads of *R. padi* head (n = 3) and body samples (n = 3) were retrieved from Thorpe, et al. 2016 through the European Nucleotide Archive under the project accession number of PRJEB9912. Sample accessions for head and body datasets are as follows: SAMEA3505144, SAMEA3505145, SAMEA3505146, SAMEA3505147, SAMEA3505148, SAMEA3505149. Reads were trimmed for adapters and quality and mapped against the reference transcriptome assembled in this study following the procedures described in the methods. Normalized expression counts (TPM) of putative salivary effectors in head and body samples were used to generate the heatmap. Each row and column represent the transcripts and biological replicates of head and body samples, respectively. Red asterisks indicate the two functionally characterized salivary effector genes. Source data is provided in the Source Data file.

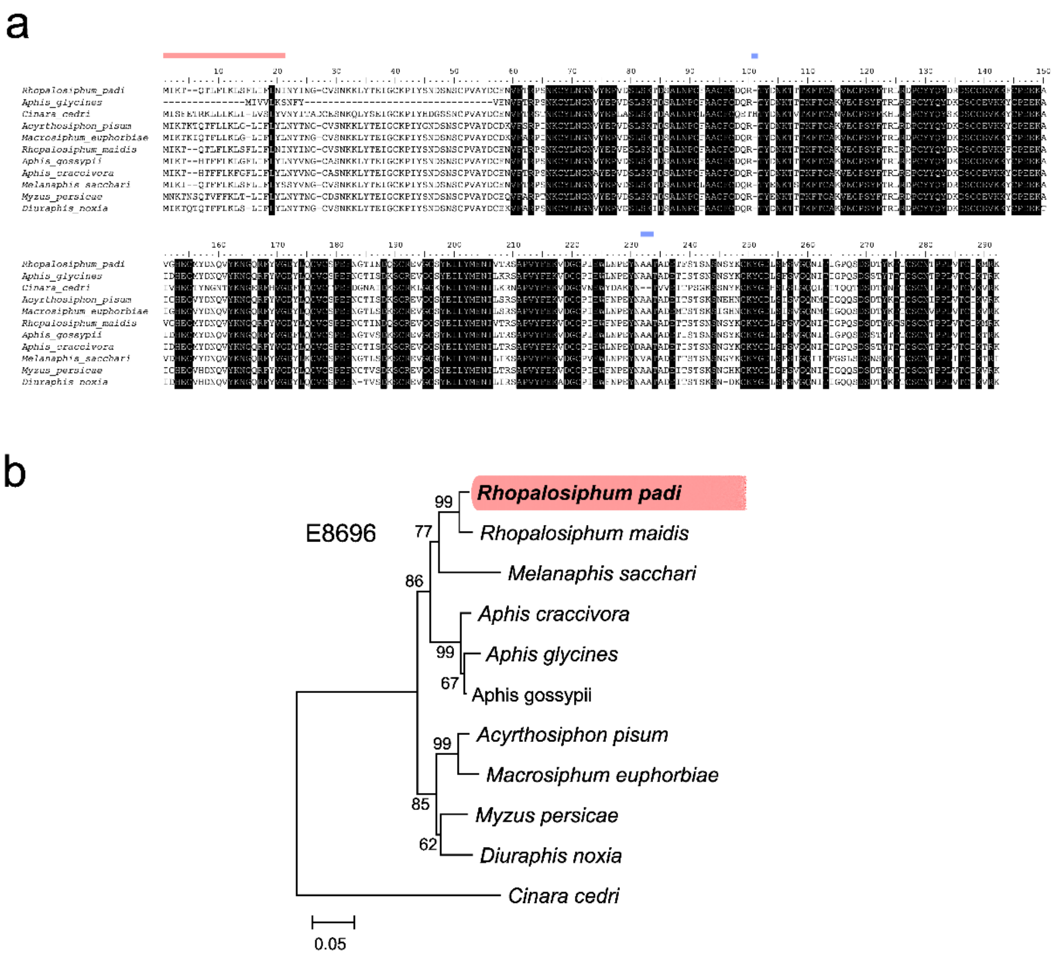


**Supplementary Figure 4. a** Multiple sequence alignment of E8696 orthologs of aphid species. Amino acid sequences of each salivary effector were aligned using Muscle. Black shade indicates the identical amino acid residues across the sequences. Pink and blue bars above the alignment depict the regions of predicted signal peptide and gaps. **b** Evolutionary relationships of E8696 across different aphid species. Phylogenetic trees were generated by the maximum likelihood method using GTR+G as substitution model. Number above the node indicates the percentage bootstrap support values over 1000 replications. Scale bar represents the number of substitutions per site. The NCBI accession numbers for homologous sequences of *R. padi* E8696 and their sequence similarities are provided in Supplementary Table 5.

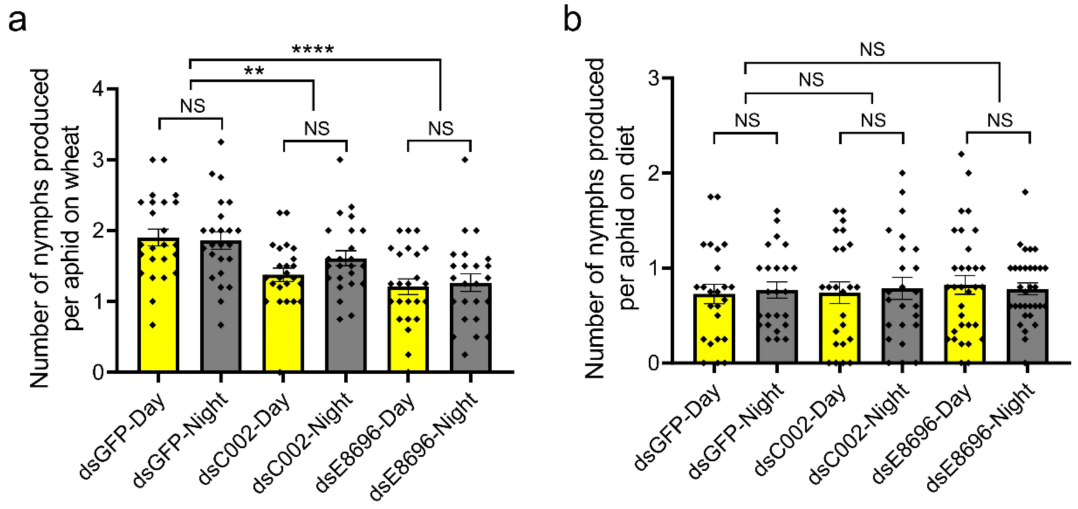


**Supplementary Figure 5.** Aphid fecundity after silencing C002 and E8696 salivary effector genes on wheat (**a**) and artificial diet (**b**). Five surviving aphids after 24h post microinjection with dsRNA were placed onto a wheat or diet. The dsRNA targeting the green fluorescent protein (GFP) gene (dsGFP) was used as negative control. Numbers of nymphs produced during the 12h daytime and 12h nighttime were counted for four days (on wheat: dsC002's n = 6, dsGFP's n = 6, dsE8696's n = 6; on diet: dsC002's n = 6, dsGFP's n = 6, dsE8696's n = 8). Each dot and bar represent the number of nymphs produced per aphid and their mean ± standard error, respectively. Statistical difference between treatments was tested using Mann-Whitney test (**adj. $P$<0.01, ****adj. $P$<0.0001). NS indicates the difference between treatments is not significant. Source data and full statistical summary are provided in the Source Data file.

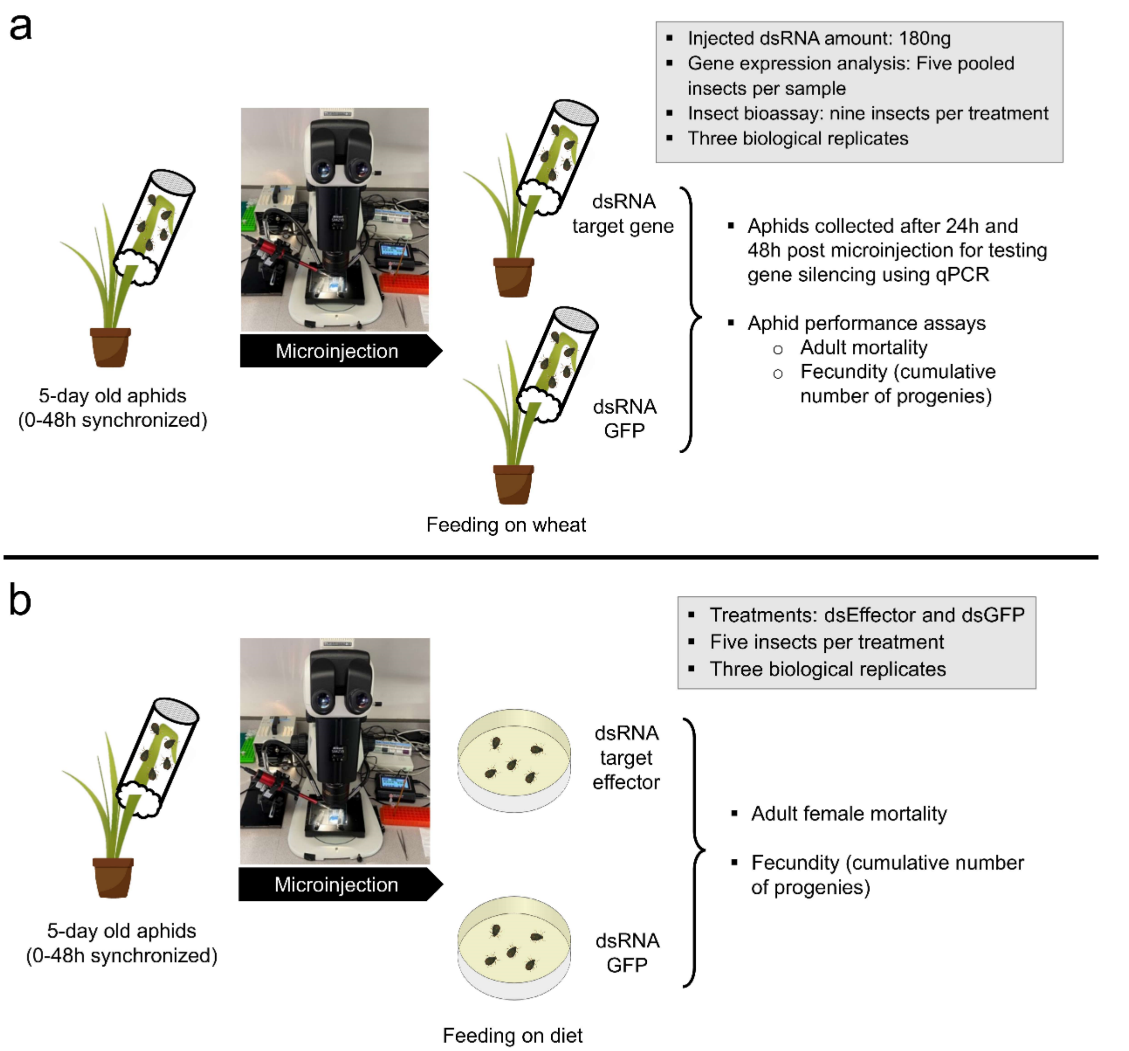


**Supplementary Figure 6.** Experimental design for microinjection of dsRNA into aphids. Aphid performance on plants (**a**) and artificial diet (**b**) was investigated after silencing salivary effector genes.

**Supplementary Table 1.** Cosinor analysis of selected aphid feeding behaviors

| Acronym | Description | Mesor[a] | Amplitude[b] | Acrophase[c] | Bathyphase[d] | Zero-amplitude test | | | |
|---|---|---|---|---|---|---|---|---|---|
| | | | | | | Df1 (ndf, numerator) | Df2 (ddf, denominator) | *F*-value | *P*-value |
| n_np | Number of np | 5.2 | 0.4 | 6.0 | 18.0 | 2 | 137 | 0.49 | 0.615 |
| s_np | Total duration of np | 1155.4 | 335.7 | 1.9 | 13.9 | 2 | 137 | 4.03 | 0.020 |
| n_Pr | Number of probes | 5.2 | 0.4 | 6.0 | 18.0 | 2 | 137 | 0.49 | 0.615 |
| s_Pr | Total probing time | 12730.0 | 254.6 | 11.5 | 23.5 | 2 | 137 | 0.32 | 0.729 |
| n_C | Number of C | 11.1 | 1.1 | 14.9 | 26.9 | 2 | 137 | 1.13 | 0.327 |
| s_C | Total duration of C | 3445.7 | 341.4 | 8.1 | 20.1 | 2 | 137 | 1.25 | 0.291 |
| n_pd | Number of pd | 41.9 | 6.8 | 11.1 | 23.1 | 2 | 137 | 2.93 | 0.057 |
| s_pd | Total duration of pd | 189.0 | 24.9 | 10.2 | 22.2 | 2 | 137 | 1.84 | 0.162 |
| n_sgE1 | Number of single E1 | 2.4 | 0.8 | 14.6 | 26.6 | 2 | 91 | 4.25 | 0.017 |
| s_sgE1 | Total duration of single E1 | 420.2 | 114.3 | 9.4 | 21.4 | 2 | 90 | 1.55 | 0.219 |
| n_E1 | Number of E1 | 5.0 | 1.9 | 12.1 | 24.1 | 2 | 135 | 9.35 | 0.000 |
| s_E1 | Total duration of E1 | 881.9 | 334.3 | 9.8 | 21.8 | 2 | 134 | 4.15 | 0.018 |
| n_E2 | Number of E2 | 2.7 | 0.6 | 12.4 | 24.4 | 2 | 135 | 4.02 | 0.020 |
| s_E2 | Total duration of E2 | 6269.0 | 174.8 | 20.9 | 32.9 | 2 | 137 | 0.08 | 0.920 |
| n_G | Number of G | 1.4 | 0.2 | 17.8 | 29.8 | 2 | 49 | 0.96 | 0.390 |
| s_G | Duration of G | 1515.3 | 482.9 | 6.1 | 18.1 | 2 | 51 | 1.69 | 0.194 |
| n_F | Number of np | 2.8 | 0.5 | 23.0 | 35.0 | 2 | 80 | 1.78 | 0.176 |
| s_F | Total duration of np | 2461.2 | 741.6 | 20.1 | 32.1 | 2 | 80 | 2.24 | 0.113 |

[a] **Mesor:** An estimate of central tendency of the distribution of values of an oscillating variable (the average value around which the variable oscillates). The mesor is a circadian rhythm-adjusted mean based on the parameters of a cosine function fitted to the raw data. Note: When a process is known to be rhythmic, and the data points are not equidistant or the sample size is small, the Mesor often provides a more appropriate unbiased estimator of central tendency than does the arithmetic mean of the raw data.

[b] **Amplitude:** The difference between the peak (or trough) and the mean value of a wave. Note: For symmetrical waves, the amplitude is half the value of the range of oscillation.

[c] **Acrophase:** The time at which the peak of a rhythm occurs. Note: Originally, acrophase referred to the phase angle of the peak of a cosine wave fitted to the raw data of a rhythm (time series). When the term is applied to the actual rhythm, the acrophase will likely vary from cycle to cycle. Unit of measurement: hours (h) or degrees of circumference (°) in relation to an absolute or arbitrary reference.

[d] **Bathyphase:** The time at which the trough of a rhythm occurs. Unit of measurement: hours (h) or degrees of circumference (°) in relation to an absolute or arbitrary reference.

**Supplementary Table 2.** Statistical summary of genome-guided transcriptome assembly for *R. padi*

| Measurement* | Number |
|---|---|
| Total assembled contigs | 49,486 |
| Total assembled bases | 109,061,758 |
| Mean length of contigs (bp) | 2,204 |
| N50 of contigs | 3,666 |
| Min/Max contig length (bp) | 200/42,258 |
| Sequences with open reading frame (ORF) | 36,932 |
| Read mapping rate to genome reference | 97% |
| BUSCO score for complete sequences | 99% |

*Quality of assembled transcriptome was evaluated using Transrate v2.14.0 and BUSCO v5.2.2 against the lineage of arthropoda_odb10. Read mapping rate was estimated by mapping reads from each library to the genome reference using Hisat2 v2.2.1.

**Supplementary Table 3.** Rhythmicity of osmoregulatory genes in *R. padi*

| Gene name | Symbol | Transcript ID | Aphidbase accession | Acrophase | Amplitude | *p*-value |
|---|---|---|---|---|---|---|
| Aquaporin 1 | *AQP1* | MSTRG.10863.9 | g15260.t1 | 15.73 | 13.98 | 0.031 |
| Gut sucrase 1 | *SUC1* | MSTRG.10130.1 | g14086.t1 | 18.09 | 11.25 | 0.013 |
| Sugar transporter | *ST* | MSTRG.10126.1 | g14083.t1 | 21.98 | 7.67 | 0.006 |

**Supplementary Table 4.** Diurnally rhythmic putative salivary effectors in *R. padi*

| Transcript ID* | Mesor | Acrophase | Amplitude | *P*-value |
|---|---|---|---|---|
| g12028.t1 | 99.9 | 18.9 | 18.5 | 0.006 |
| g12266.t1 | 411.8 | 18.0 | 76.1 | 0.008 |
| g12327.t1 | 92.8 | 6.9 | 29.2 | 0.006 |
| g13874.t1 | 191.3 | 22.3 | 20.8 | 0.032 |
| g14344.t1 | 26.6 | 2.5 | 7.1 | 0.009 |
| g147.t1 | 109.8 | 8.8 | 31.0 | 0.047 |
| g14944.t1 | 12.7 | 3.0 | 2.0 | 0.045 |
| g14946.t1 | 32.2 | 4.5 | 5.0 | 0.049 |
| g15504.t1 | 352.2 | 7.0 | 78.0 | 0.030 |
| g15702.t1 | 304.0 | 7.9 | 74.5 | 0.039 |
| g18650.t1 | 408.1 | 7.2 | 166.4 | 0.001 |
| g19280.t1 | 43.2 | 5.0 | 39.8 | 0.016 |
| g21517.t1 | 120.8 | 7.9 | 31.6 | 0.004 |
| g218.t1 | 7.3 | 5.1 | 1.6 | 0.017 |
| g3181.t1 | 40.4 | 6.3 | 16.1 | 0.000 |
| g3188.t1 | 21.2 | 7.5 | 8.3 | 0.000 |
| g3960.t1 | 280.9 | 18.5 | 50.0 | 0.004 |
| g478.t1 | 25.1 | 7.1 | 9.2 | 0.026 |
| g55.t1 | 154.9 | 7.6 | 40.1 | 0.017 |
| g5983.t1 | 383.4 | 4.4 | 63.8 | 0.005 |
| g6826.t1 | 439.0 | 18.2 | 87.4 | 0.002 |
| g817.t1 | 214.6 | 18.2 | 46.2 | 0.028 |
| g8251.t1 | 44.0 | 8.3 | 17.8 | 0.015 |
| g8696.t1 | 241.9 | 17.7 | 114.6 | 0.001 |
| MSTRG.10669.1 | 11.3 | 4.0 | 1.3 | 0.049 |
| MSTRG.10699.6 | 7.4 | 2.1 | 1.4 | 0.031 |
| MSTRG.11013.1 | 157.8 | 6.5 | 38.6 | 0.007 |
| MSTRG.11258.1 | 23.4 | 4.6 | 4.8 | 0.012 |
| MSTRG.11258.2 | 31.2 | 6.0 | 7.7 | 0.001 |
| MSTRG.1199.1 | 68.2 | 23.6 | 13.5 | 0.034 |
| MSTRG.13338.2 | 26.0 | 3.8 | 7.8 | 0.002 |
| MSTRG.15390.10 | 0.6 | 22.6 | 0.4 | 0.013 |
| MSTRG.15390.2 | 15.0 | 18.1 | 4.8 | 0.018 |
| MSTRG.1801.2 | 32.1 | 8.0 | 6.7 | 0.024 |
| MSTRG.1801.4 | 13.1 | 6.5 | 3.6 | 0.010 |
| MSTRG.2389.1 | 18.5 | 6.5 | 6.8 | 0.000 |
| MSTRG.3386.2 | 0.2 | 19.4 | 0.4 | 0.044 |
| MSTRG.442.4 | 6.1 | 19.1 | 1.4 | 0.045 |
| MSTRG.5122.3 | 12.7 | 22.1 | 3.7 | 0.015 |
| MSTRG.5899.4 | 0.8 | 9.8 | 0.7 | 0.024 |
| MSTRG.8529.1 | 123.6 | 5.8 | 26.5 | 0.006 |
| MSTRG.8802.2 | 3.1 | 22.4 | 1.3 | 0.013 |

*Among a total of 264 putative salivary effector transcripts, 42 were identified as diurnally rhythmic.

**Supplementary Table 5.** Identification of *R. padi* E8696 homologs in aphid species

| Species | NCBI mRNA accession | NCBI protein accession | Identity % | Coverage % | *E*-value |
|---|---|---|---|---|---|
| *Aphis glycines* | VYZN01000065.1 | KAE9524594.1 | 94.44 | 80 | 5E-165 |
| *Cinara cedri* | CABPRJ010000950.1 | VVC31548.1 | 69.49 | 93 | 3E-138 |
| *Acyrthosiphon pisum* | XM_029491156.1 | XP_029347016.1 | 88.81 | 98 | 0 |
| *Macrosiphum euphorbiae* | CARXXK010000004.1 | CAI6366975.1 | 88.28 | 99 | 0 |
| *Rhopalosiphum maidis* | XM_026949305.2 | XP_026805106.1 | 98.26 | 99 | 0 |
| *Aphis gossypii* | XM_027989221.2 | XP_027845022.1 | 92.71 | 99 | 0 |
| *Aphis craccivora* | VUJU01000964.1 | KAF0767456.1 | 92.01 | 99 | 0 |
| *Melanaphis sacchari* | XM_025345618.1 | XP_025201403.1 | 88.85 | 99 | 0 |
| *Myzus persicae* | XM_022312603.1 | XP_022168295.1 | 89.47 | 98 | 0 |
| *Diuraphis noxia* | XM_015519836.1 | XP_015375322.1 | 88.97 | 99 | 0 |

**Supplementary Table 6.** Primers used for dsRNA synthesis and gene expression analysis

| Gene | Primer name | Strand | Sequence (5' to 3')* | Length | Annealing temperature | Amplicon size |
|---|---|---|---|---|---|---|
| AQP1 | AQP1-T7F | F | GGATCCTAATACGACTCACTATAGGCGGATTAGGTTCATCGGCTA | 45 | - | - |
| | AQP1-T7R | R | GGATCCTAATACGACTCACTATAGGGGTTTGTCCAAAAGCCAAGA | 45 | - | |
| | AQP1-F | F | CGGATTAGGTTCATCGGCTA | 20 | 60 | 500 |
| | AQP1-R | R | GGTTTGTCCAAAAGCCAAGA | 20 | 60 | |
| | AQP1-RTF | F | GATCTTTAGGCCCAGCCGTT | 20 | 60 | 100 |
| | AQP1-RTR | R | GTGTGGATGGTTCCTCCAAGT | 21 | 60 | |
| SUC1 | SUC1-T7F | F | GGATCCTAATACGACTCACTATAGGCGATTCTTTGACGCATACGA | 45 | - | - |
| | SUC1-T7R | R | GGATCCTAATACGACTCACTATAGGCCCTGCTGGATCAACAGTTT | 45 | - | |
| | SUC1-F | F | CGATTCTTTGACGCATACGA | 20 | 60 | 478 |
| | SUC1-R | R | CCCTGCTGGATCAACAGTTT | 20 | 60 | |
| | SUC1-RTF | F | GGATCTGCGTGGGAATGGAA | 20 | 60 | 156 |
| | SUC1-RTR | R | TCTAAACCCATCGACGCCAC | 20 | 60 | |
| C002 | C002-T7F | F | TAATACGACTCACTATAGGGAGAGCTAGTGCTGCTGACGTGTA | 45 | - | - |
| | C002-T7R | R | TAATACGACTCACTATAGGGAGAACGTGACGTCTACCTCTCTCA | 45 | - | |
| | C002-F | F | GCTAGTGCTGCTGACGTGTA | 20 | 60 | 484 |
| | C002-R | R | ACGTGACGTCTACCTCTCTCA | 20 | 60 | |
| | C002-RTF | F | AGTTCGTAGAGACAAAGGGCG | 21 | 57 | 101 |
| | C002-RTR | R | CTTGAGCAAGTTGATGACCCG | 21 | 56 | |
| E8696 | E8696-T7F | F | GGATCCTAATACGACTCACTATAGGACATGTGCCAAGGTTGAATG | 45 | - | - |
| | E8696-T7R | R | GGATCCTAATACGACTCACTATAGGGAGTCTGACTGTGGGCCAAT | 45 | - | |
| | E8696-F | F | ACATGTGCCAAGGTTGAATG | 20 | 59 | 476 |
| | E8696-R | R | GAGTCTGACTGTGGGCCAAT | 20 | 60 | |

| | | | | | | |
|---|---|---|---|---|---|---|
| | E8696-RTF | F | ACCCATGTTTTGCTGCCTGT | 20 | 60 | 90 |
| | E8696-RTR | R | ATGGACATTCAACCTTGGCAC | 21 | 59 | |
| GFP | GFP-T7F | F | GGATCCTAATACGACTCACTATAGGGGC ACGACTTCTTCAAGAGC | 45 | - | - |
| | GFP-T7R | R | GGATCCTAATACGACTCACTATAGGCAT GCCATGTGTAATCCCAG | 45 | - | |
| | GFP-F | F | GGCACGACTTCTTCAAGAGC | 20 | 60 | 461 |
| | GFP-R | R | CATGCCATGTGTAATCCCAG | 20 | 60 | |
| Actin | Actin-RTF | F | TGGTATTGCCGACAGAATGC | 20 | 58 | 189 |
| | Actin-RTR | R | ACGATGGATGGGCCTGATTC | 20 | 59 | |

**Supplementary Data 1 (Separate file).** Results for Electrical penetration graph (EPG) parameters measured for aphids feeding on wheat plants over 24 hours

**Supplementary Data 2 (Separate file).** Complete list of diurnally rhythmic transcripts in the whole-body *R. padi*

**Supplementary Data 3 (Separate file).** Clustering and annotation of diurnally rhythmic transcripts in the whole-body *R. padi*

**Supplementary Data 4 (Separate file).** *In silico* identification and annotation of putative salivary effectors in *R. padi*

**Supplementary Data 5 (Separate file).** Results for Electrical penetration graph (EPG) parameters measured for aphids feeding on wheat plants after silencing E8696 salivary effector gene
